# TNF stimulation primarily modulates transcriptional burst size of NF-κB-regulated genes

**DOI:** 10.1101/2020.11.16.384297

**Authors:** Victor L. Bass, Victor C. Wong, M. Elise Bullock, Suzanne Gaudet, Kathryn Miller-Jensen

## Abstract

Cell-to-cell heterogeneity is a characteristic feature of the tumor necrosis factor (TNF)-stimulated inflammatory response mediated by the transcription factor NF-κB, motivating an exploration of the underlying sources of this noise. Here we combined single-transcript measurements with computational models to study transcriptional noise at six NF-κB-regulated inflammatory genes. In the basal state, NF-κB-target genes displayed an inverse correlation between mean and noise. TNF stimulation increased transcription while maintaining noise, except for the most repressed genes. By fitting transcript distributions to a two-state model of promoter activity, we found that TNF primarily stimulated transcription by increasing burst size while maintaining burst frequency. Burst size increases were associated with enrichment of initiated-but-paused RNA polymerase II at the promoter, and blocking the release of paused RNAPII with a small molecule inhibitor decreased TNF-stimulated burst size. Finally, we used a mathematical model to show that TNF positive feedback further amplified gene expression noise resulting from burst-size mediated transcription, leading to diverse TNF functional outputs. Our results reveal potential sources of noise underlying intercellular heterogeneity in the TNF-mediated inflammatory response.

## Introduction

Tumor necrosis factor (TNF) activates pro-inflammatory and stress response signaling in many cell types (Aggarwal, 2003). The TNF inflammatory response is mediated by the transcription factor NF-κB, which regulates the expression of hundreds of genes. These genes include inflammatory cytokines that can propagate an immune response via paracrine signaling, as well as negative regulators of NF-κB (Hoffmann *et al*, 2002; Pahl, 1999; Smale, 2011). Dysregulation of the TNF-stimulated NF-κB response contributes to inflammatory disease states (Lewis and Pollard, 2006; Schottenfeld and Beebe-Dimmer, 2006.), and thus NF-κB-induced transcription is tightly regulated in cell populations. However, it has been widely observed that TNF stimulates significant cell-to-cell heterogeneity in NF-κB signaling and in the transcription of its inflammatory gene targets (Cheong *et al*, 2011; Lee *et al*, 2014; Tay *et al*, 2010; Wong *et al*, 2019; Zhang *et al*, 2017). Although cell-to-cell heterogeneity in NF-κB signaling has been widely explored, additional sources of noise underlying transcription are not well understood. Understanding these sources of noise may enhance our ability to modulate the inflammatory response in clinically relevant ways.

One major source of single-cell gene expression noise is the fluctuation of promoters between transcriptionally active and inactive states, a process termed transcriptional bursting (Bahar Halpern *et al*, 2015b; Dar *et al*, 2012; Raj *et al*, 2006. Singh *et al*, 2010; Skupsky *et al*, 2010; Suter *et al*, 2011.). Though gene expression noise can be buffered by various mechanisms (Bahar Halpern *et al*, 2015a; Padovan-Merhar *et al*, 2015; Stoeger *et al*, 2016), in some cases it is amplified by regulatory networks to drive diverse cellular behaviors (Acar *et al*, 2008; Chang *et al*, 2008; Shalek *et al*, 2014; Weinberger *et al*, 2005). Several molecular mechanisms have been associated with transcriptional bursting including nucleosome positioning (Dey *et al*, 2015; Raser and O’Shea, 2004), chromatin modifications (Chen *et al*, 2019; Suter *et al*, 2011), transcription factor activity (Li *et al*, 2018; Senecal *et al*, 2014), and RNA polymerase (RNAPII) pause regulation (Bartman *et al*, 2019; Wong *et al*, 2018).

Although transcriptional bursting has not been extensively studied at endogenous NF-κB target genes, it has been well characterized for the HIV long terminal repeat (LTR) promoter, which is regulated by NF-κB. Transcriptional bursting at the HIV LTR has been shown to be influenced by chromatin environment both in the basal state (Dey *et al*, 2015; Dar *et al*, 2012; Singh *et al*, 2010), and after TNF stimulation (Dar *et al*, 2012; Wong *et al*, 2018). Specifically, it was shown that TNF could modulate either burst frequency (i.e., the rate of transition from an inactive to active state promoter state), or burst size (i.e., the number of transcripts produced per burst) of silent-but-inducible HIV LTR promoters, and that the bursting mechanism was influenced by the basal histone 3 acetylation state at the promoter (Wong *et al*, 2018). Endogenous NF-κB target promoters are found in basal chromatin environments that resemble those of latent-but-inducible HIV promoters (Ramirez-Carrozzi *et al*, 2009). Thus, we sought to determine if molecular mechanisms regulating transcriptional bursting at inducible HIV LTRs are similar for endogenous NF-κB targets.

In this study, we analyzed changes in gene expression noise and transcriptional bursting at six endogenous NF-κB target promoters before and after TNF stimulation. We found that NF-κB-target genes display an inverse correlation between mean and noise in the basal state, and that TNF stimulation increases mean transcription while maintaining noise for all but the most repressed genes. Using a mathematical model of bursting (Raj *et al*, 2006; Dey *et al*, 2015; Wong *et al*, 2018), we inferred that TNF stimulation primarily increases burst size while maintaining burst frequency, leading to highly skewed transcript distributions, especially for *Tnf* and *Il8*. We further found that differences in RNA polymerase (RNAPII) pause regulation are associated with differences in the regulation of transcriptional bursting in response to TNF, and can be modulated by altering RNAPII pause regulation with a small molecule inhibitor. Finally, we used a mathematical model to explore how TNF positive feedback further affects cell-to-cell heterogeneity in *Tnf* transcription when activated by burst size vs. burst frequency increases. We find that burst size increases, when amplified by positive feedback, lead to significantly more heterogeneous cell populations, with a small subset of high TNF producers, consistent with observations from other studies. Overall, we conclude that TNF-mediated transcriptional bursting is regulated similarly for endogenous and viral NF-κB target promoters. Moreover, our results suggest that burst size-mediated transcription combined with positive feedback may contribute to the substantial cell-to-cell variability observed in the TNF-mediated inflammatory response.

## Results

### Single-molecule mRNA quantification reveals a conserved mean-noise relationship for TNF-NF-κB gene targets in the basal state

To characterize transcriptional noise in NF-κB targets induced by TNF, we analyzed six genes regulated by NF-κB. These genes have different roles in the TNF-induced inflammatory response. *Nfkbia* and *Tnfaip3* encode the intracellular proteins IκB-α and A20, respectively, which negatively regulate NF-κB p65 (Baeurerle and Baltimore, 1988; Heyniinck *et al*, 1999; Hoffmann *et al*, 2002). *Tnf, Il8, Il6* and *Csf2* encode the secreted inflammatory cytokines TNF, IL-8, IL-6, and GM-CSF, respectively (Fig. 1A). *Nfkbia, Tnfaip3, Tnf*, and *Il8* are classified as primary inflammatory genes because they are transcribed directly in response to stimulation in immune cells, while *Il6* and *Csf2* are classified as secondary genes because they require synthesis of additional protein regulators prior to transcription (Hargreaves *et al*, 2009; Ramirez-Carrozzi *et al*, 2006; Ramirez-Carrozzi *et al*, 2009). These genes are found in a range of basal chromatin environments, as quantified by the ratio of histone H3 acetylated at lysine 9 and 14 (AcH3) to total histone H3 levels (AcH3:H3) at their promoters measured by chromatin immunoprecipitation (ChIP) in the leukemic Jurkat T cell line. *Nfkbia* and *Tnfaip3* had the highest ratio of AcH3:H3, indicating a more open chromatin environment, while *Il6* and *Csf2* had much lower ratios, indicating a more closed chromatin state (Fig. 1B).

**Figure 1.**
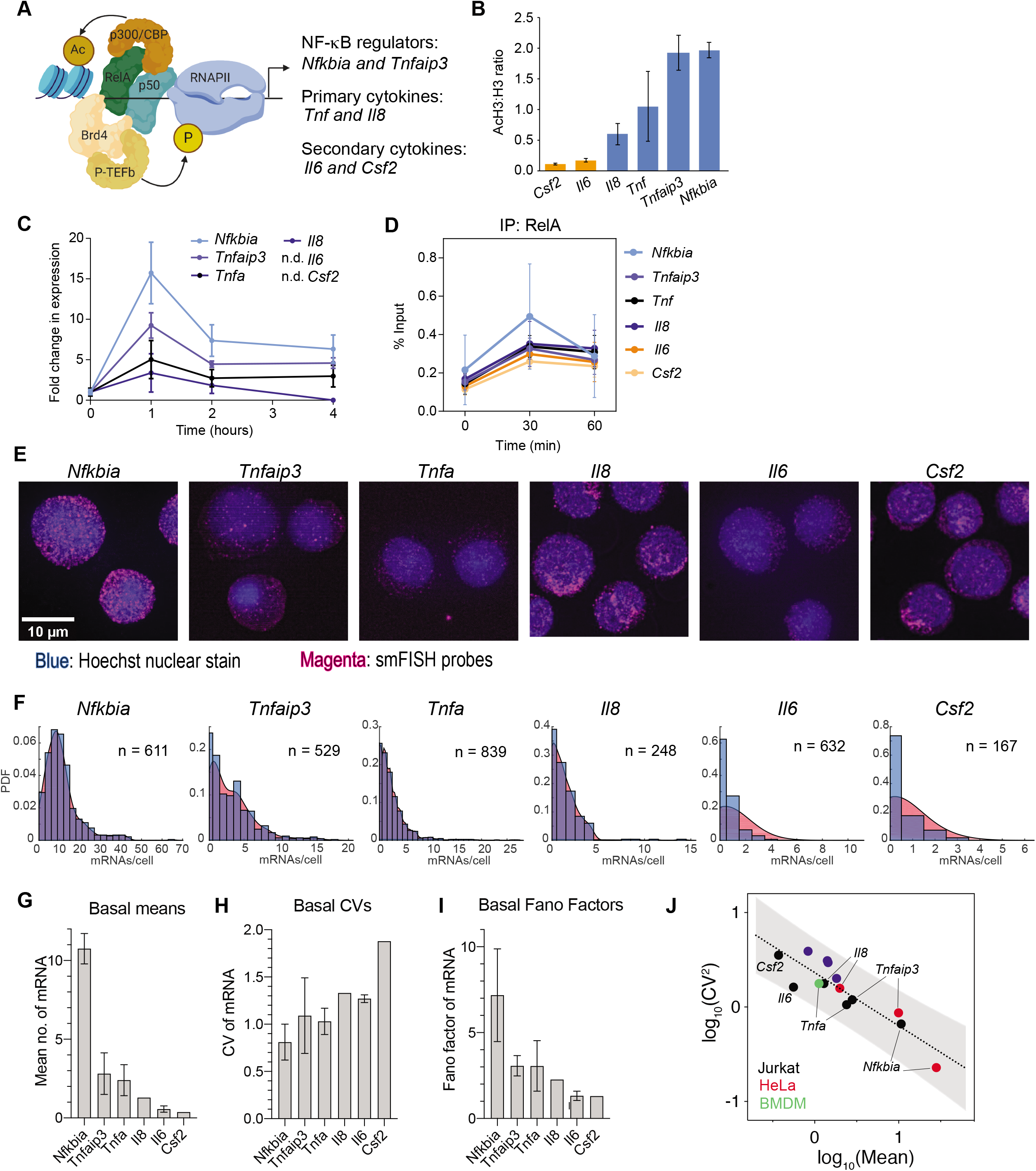
Basal transcriptional noise of NF-κB targets is systematically related to basal chromatin environment. **A** NF-κB can recruit a variety of binding partners to target promoters, including the chromatin modifying enzyme p300, the elongation complex P-TEFb, and components of transcriptional machinery. NF-κB target genes with a variety of functions were chosen for this study. **B** Ratio of enrichment of total histone H3 to acetylated H3 (AcH3) in Jurkat T cells for the indicated targets quantified by ChIP-qPCR and shown as % input (non-IP control). Data are presented as mean ± standard deviation (SD) of three biological replicates. **C** Induction of NF-κB targets in Jurkat T cells in response to 20 ng/ml TNF treatment for 1, 2, and 4 hours as measured by RT-qPCR. Target values were normalized to GAPDH and are shown as fold change relative to basal expression is shown as mean ± SD of 3 biological replicates. **D** Enrichment of RelA before and 30 and 60 minutes after treatment with 20 ng/mL TNF as measured by ChIP-qPCR and shown as % input (non-IP control). Data are presented as mean ± SD of three biological replicates. **E** Maximum intensity projections of smFISH fluorescence microscopy z-stacks of basal Jurkat T cells stained for the indicated genes. Images were filtered with a dual Gaussian filter in the FISH-quant software program and then brightness and contrast enhanced for better visualization. Scale bars: 10 μM. **F** Histograms of transcripts per cell for target genes (blue) overlaid with probability density plots (red) generated from smFISH data. **G-I** Bar graphs of mean (G), CV (H), and Fano (I) of smFISH distributions for the indicated genes. Data are presented as mean ± bootstrapped 95% confidence intervals (CIs). **J** Log-log graph of mean versus noise (CV^2^) of basal mRNA distributions measured in Jurkat T cells (black), HeLa cells (red) or murine bone marrow derived macrophages (green) for endogenous genes and four HIT lentivirus integrations in Jurkats (blue). Gray shading indicates 95% CI of the linear regression for the basal burst size trend line. HeLa data from Lee *et al*. (2014) and LTR data from Wong *et al*. (2018).

In response to TNF stimulation (20 ng/ml), most genes exhibited increased transcription that inversely correlated with AcH3:H3 ratio in the basal state: *Nfkbia* and *Tnfaip3* exhibited the highest increases, while *Tnf* and *Il8* were significantly lower, as measured in the population by RT-qPCR (Fig. 1C). Increases in *Il6* and *Csf2* were not detectable in the population even four hours after TNF stimulation. Notably, the differences in transcription were not due to differences in NF-κB p65 binding, because following TNF stimulation, NF-κB p65 promoter binding increased similarly across all promoters as measured by ChIP, including at the *Il6* and *Csf2* promoters (Fig. 1D).

To quantify transcription in single cells, we performed single molecule RNA fluorescence in situ hybridization (smFISH) in Jurkat T cells (Fig. 1E) (Raj *et al*, 2008). We found very low levels of basal transcription, ranging from an average of 10 mRNAs per cell for *Nfkbia* to less than one mRNA on average per cell for *Il6* and *Csf2* (Fig. 1F-G). Average mRNA levels were inversely correlated with AcH3:H3 ratio. We observed significant cell-to-cell heterogeneity as measured by coefficient of variation (CV) and Fano factor, with higher CV for the lower expression genes and higher Fano for the higher expression genes (Fig. 1H-I). Thus, the influence of chromatin state is apparent in the mean and variability of basal mRNA levels prior to TNF stimulation.

There is evidence for global constraints on transcriptional noise in mammalian cells (Sanchez and Golding, 2013), and our observation of systematic changes in mean and noise across NF-κB targets in different chromatin environments is consistent with this hypothesis. To explore this further, we plotted the log of mean (log_10_(mean)) versus the log of noise (log_10_(CV^2^)) for the basal mRNA measurements of these six NF-κB-regulated genes in Jurkat cells, and we observed an inverse linear relationship between noise and mean (Fig. 1J). Interestingly, when we plotted log_10_(mean) versus log_10_(CV^2^) for smFISH measurements of the same targets made in HeLa cells (Lee *et al*, 2014) or in murine bone marrow-derived macrophage cells, we found that these measurements fell along the same line. Furthermore, basal mRNA measurements for exogenous HIV-LTR promoters measured in Jurkat T cells and exhibiting similar basal chromatin states (Wong *et al*, 2018) also fell along the same trendline (R^2^ = 0.79). The slope of this conserved mean-versus-noise trendline suggests non-Poissonian stochastic transcription rather than continuous transcription in inducible NF-κB targets (Dar *et al*, 2016; Singh *et al*, 2010). Altogether we conclude that there is an inverse relationship between mean and noise in the basal state that is conserved across NF-κB targets and multiple cell types.

### TNF stimulation produces gene-specific changes in transcriptional heterogeneity

Genes in such diverse basal environments likely require recruitment of different factors by NF-κB to effectively activate transcription, which may lead to systematic differences in single-cell transcription distributions following stimulation (Neuert *et al*, 2013; Senecal *et al*, 2014). To analyze how transcriptional noise is altered by TNF-induced activation of NF-κB, we quantified mRNA using smFISH (Fig. 2A). For *Nfkbia, Tnfaip3, Tnf*, and *Il8*, we measured mRNA counts at one- and two-hours post TNF treatment to capture the peak and reduction in expression (Fig. 2B and Fig. 1C). For *Il6* and *Csf2*, we measured mRNA counts at 2- and 4-hours post TNF treatment when transcription was still rising. Notably, we were able to measure a significant increase in mRNA levels for *Il6* and *Csf2* by smFISH, even though increases in transcription were not detectable by population-level RT-qPCR (Fig. 1C).

**Figure 2.**
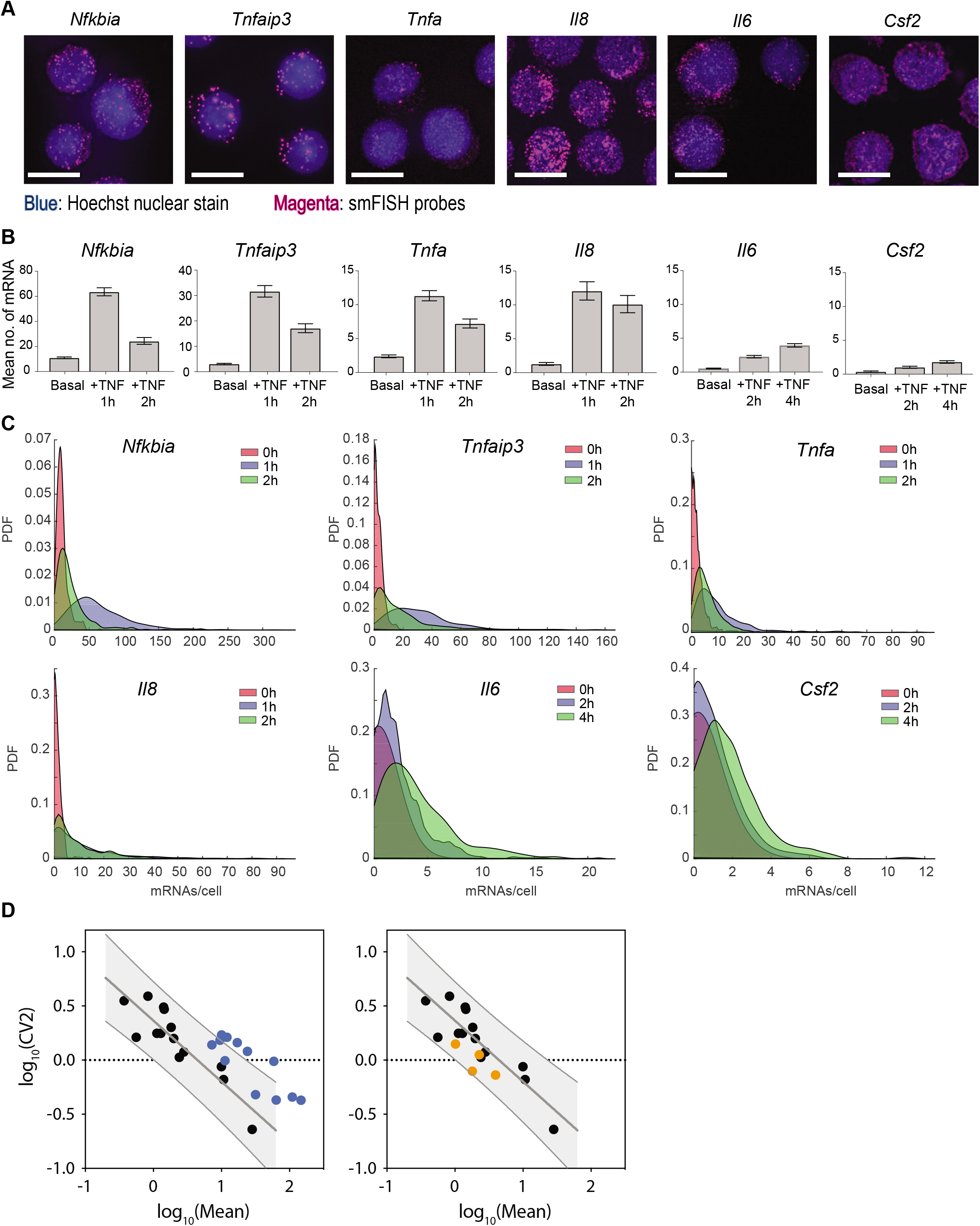
TNF induces gene-specific changes in transcript distributions at NF-κB targets. **A** Maximum intensity projections of smFISH fluorescence microscopy z-stacks of basal Jurkat T cells stained for the indicated genes after treatment with 20 ng/mL TNF. Images were filtered with a dual gaussian filter in the FISH-quant software program and brightness and contrast enhanced for better visualization. Scale bars: 10 μM. **B** Bar graphs of mean of smFISH distributions before and after TNF treatment for the indicated genes. Data are presented as mean ± bootstrapped 95% CIs. **C** Probability density plots of single cell mRNA distributions from smFISH before and after treatment with 20 ng/mL TNF for the indicated time points. **D** Log-log graph of mean versus noise showing shifts in transcript distributions after treatment with 20ng/mL TNF grouped by whether noise and mean increase together (left) or noise decreases as mean increases (right). Gray shading indicates 95% CI of basal trend.

Although TNF treatment increased mean mRNA counts for all targets, the change in transcriptional noise varied by gene, as observed from the single-cell mRNA distributions (Fig. C). After TNF treatment, *Nfkbia* and *Tnfaip3* were expressed in most cells, while *Tnf* and *Il8* were expressed at lower levels with more non-expressing cells, but all four targets exhibited long-tailed distributions, with a few cells expressing mRNA counts much higher than the mean. In contrast, *Il6* and *Csf2* were expressed at much lower levels and exhibited less skewed distributions (Fig. 2B).

These differences were apparent when observing the dynamic trends in CV and Fano factor. For *Nfkiba, Tnfaip3, Tnf*, and *Il8*, the CV of mRNA counts remained relatively constant from 0-2 hours, while the CV of mRNA counts for *Il6* and *Csf2* decreased from 0-4 hours (Fig. EV1A). In contrast, the Fano factor for *Nfkiba*, *Tnfaip3*, *Tnf*, and *Il8* increased significantly over time, while the Fano factor for *Il6* and *Csf2* remained relatively constant (Fig. EV1B). We found no significant reductions in noise after normalizing for cell area (Fig. EV1C), in contrast to other targets for which single-cell mRNA expression has been shown to correlate with cell size (Bagnall *et al*, 2018; Padovan-Merhar *et al*, 2015). This lack of correlation with cell size suggests that, for these inflammatory gene targets, shared sources of cellular variation are less important than gene-specific noise sources.

To visualize how TNF-NF-κB-mediated transcription changed the global mean-noise relationship seen in the basal state, we plotted log_10_(mean) and log_10_(CV^2^) of mRNA counts before and after TNF treatment. We found that for the NF-κB targets that increased mean without a significant reduction in noise, we observed that in some cases they moved outside the basal trendline, especially at 2 hours (Fig. 2D, left). In contrast, *Il6* and *Csf2* remained within the trendline of the basal measurements upon TNF treatment for 2 and 4 hours as noise decreased with increased mean (Fig. 2D, right). Overall, this suggests that NF-κB differentially regulates transcriptional noise at different target genes following TNF stimulation.

### TNF stimulation primarily modulates burst size of NF-κB targets

For many mammalian genes, transcription occurs in short bursts. Bursting behavior can be effectively modeled with two promoter states, in which a promoter briefly switches from an ‘off’ state to a transcript-producing ‘on’ state, before switching back to the ‘off’ state (Fig. 3A) (Bahar Halpern *et al*, 2015b; Dar *et al*, 2012; Raj *et al*, 2006; Singh *et al*, 2010; Skupsky *et al*, 2010; Suter *et al*, 2011). In this model, increasing transcription by increasing the frequency of bursts results in a lower CV, while increasing transcription by increasing the size of transcriptional bursts results in a higher Fano factor. Interpreted in this way, the mean-versus-noise plots suggest TNF differentially regulates burst frequency and burst size across different NF-κB target genes (Fig. 2D) (Sanchez and Golding, 2013). Therefore, we hypothesized that TNF-stimulated transcript distributions might be well described by the two-state bursting model of transcription, and that this model might provide insight into the observed differences in transcriptional noise across NF-κB target genes.

**Figure 3.**
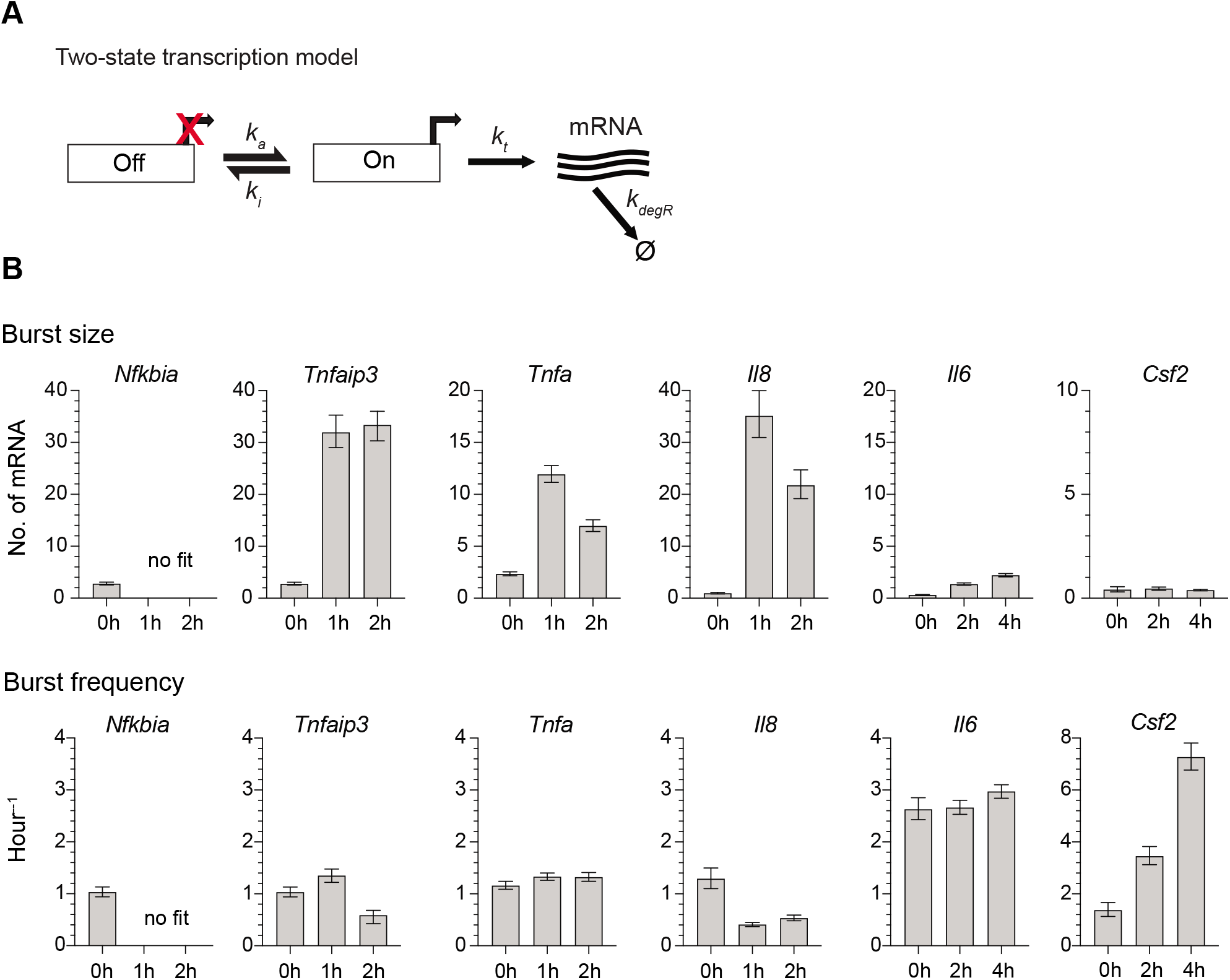
Inferred fits from two-state promoter model shows that TNF treatment increases transcriptional burst size at most targets. **A** Schematic of a two-state promoter model for transcriptional bursting. Burst frequency (k_a_) and burst size (b = k_t_/k_i_) were fit based on transcript distributions measured via smFISH. **B** Burst size (top) and burst frequency (bottom) parameter fits from the two-state model in the basal state and after treatment with 20 ng/mL TNF for 1, 2 or 4 hours. Data are presented as mean ± bootstrapped 95% CIs.

To test this, we fit our data to a two-state model of promoter activity (Raj *et al*, 2006; Dey *et al*, 2015) (Fig. 3A). In this model, also known as the random telegraph model, transcription is described by four parameters: rate of transition to the active state, k_a_; rate of transition to the inactive state, k_i_; rate of transcription in the active state, k_t_; and mRNA degradation rate, k_deg_. The probability density function (pdf) of this distribution can be solved theoretically and then burst frequency (k_a_) and burst size (mean number of transcripts produce per active state burst, b = k_t_/k_i_) can be inferred by finding the optimum fit between the experimental and theoretical pdfs using maximum likelihood estimation (MLE) (Raj *et al*, 2006; Dey *et al*, 2015; Wong *et al*, 2018).

To perform MLE, we fixed mRNA decay rate (k_deg_) to experimentally measured values when possible (Fig. EV2A and Methods). Transcription of *Il6* and *Csf2* was too low to be measured accurately, and so we used the average decay rate measured for the other four targets, which displayed similar transcript stability (t_1/2_ ≈ 40 minutes) and is in line with previously reported values (Paschoud *et al*, 2006). Because burst size is a ratio of k_t_ to k_i_, we fixed k_t_ and used MLE to infer k_i_ as previously described (Dey *et al*, 2015). For each gene, we performed a sensitivity analysis for how estimations of burst size and burst frequency varied with changes in the production rate (k_t_; Fig. EV2B). We chose values of k_t_ within a relatively insensitive range of burst size values.

After fixing k_t_ and k_deg_, we fit our single-cell transcript distributions before and after TNF treatment to the two-state model’s pdf and found that the model fit all basal distributions and most TNF-stimulated distributions (Fig. EV4). The one exception was for *Nfkbia*, in which the theoretical pdf from the two-state model could not be accurately fit to the TNF-stimulated distributions. This is consistent with the high transcriptional rate induced by TNF that might be outside the limits of the two-state bursting model (Wilson *et al*, 2017). The model fits indicate that in the basal state, most genes share a low basal burst frequency of ~1 transition per hour and a burst size of only a few transcripts (Fig. 3B). TNF treatment drives large increases in burst size with minimal changes in burst frequency for *Tnfaip3*, *Tnf*, and *Il8*. In contrast, TNF causes a small increase in both burst size and frequency for *Il6*, and a large increase in burst frequency with no change in burst size for *Csf2* (Fig. 3B). This suggests that TNF stimulation primarily alters the burst size of promoters that exhibit relatively open chromatin environments in the basal state. In contrast, for *Il6* and *Csf2*, which exhibit more closed chromatin in the basal state, TNF stimulation only modestly affects burst size or, in the case of *Csf2*, modulates burst frequency. These results are consistent with our observations at HIV-LTRs integrated in different chromatin environments (Wong *et al*, 2018) and suggest that mechanisms of transcriptional bursting are affected by the chromatin state at the promoter.

### TNF-mediated increases in burst size are associated with RNAPII pausing

Activation of a range of transcription factors (TFs) has been associated with changes in burst frequency for many genes (Chen *et al*, 2019; Friedrich *et al*, 2019; Li *et al*, 2018), but TF-mediated changes in burst size have not been as widely reported. To search for potential differences in molecular events linked to changes in burst size after TNF treatment, we measured chromatin features and binding of transcriptional machinery at our target promoters using ChIP. Changes in transcriptional burst frequency have been linked to histone acetylation (Chen *et al*, 2019; Nicolas *et al*, 2018), and we previously showed that TNF-NF-κB-mediated increases in burst size at the HIV LTR were associated with regulation of RNAPII activity (Wong *et al*, 2018). Therefore, we focused on measuring histone H3 acetylation and markers of RNAPII regulation.

Specifically, we measured total RNAPII, serine-5 phosphorylated RNAPII (initiating), serine-2 phosphorylated RNAPII (elongating), and negative elongation factor (NELF) before and at 2 and 4 hours after TNF treatment (Fig. 4A). We found that *Il6* and *Csf2* accumulated less total RNAPII than *Nfkbia, Tnfaip3, Tnf*, and *Il8*, which is consistent with the lower relative expression levels of these genes after TNF treatment. The disparity in RNAPII enrichment was lessened when looking at ser2-p RNAPII (associated with elongation) and heightened when looking at ser5-p RNAPII (associated with initiation). Enrichment of NELF, which inhibits elongation, coupled with enrichment of ser5-p RNAPII is indicative of paused RNAPII at *Tnfaip3, Tnf*, and *Il8*, in contrast to the *Il6* and *Csf2* promoters. Taken together, the RNAPII ChIP shows that the *Tnfaip3, Tnf*, and *Il8* promoters, which increase burst size after TNF treatment, accumulate more paused RNAPII than the *Il6* and *Csf2* promoters in response to TNF.

**Figure 4.**
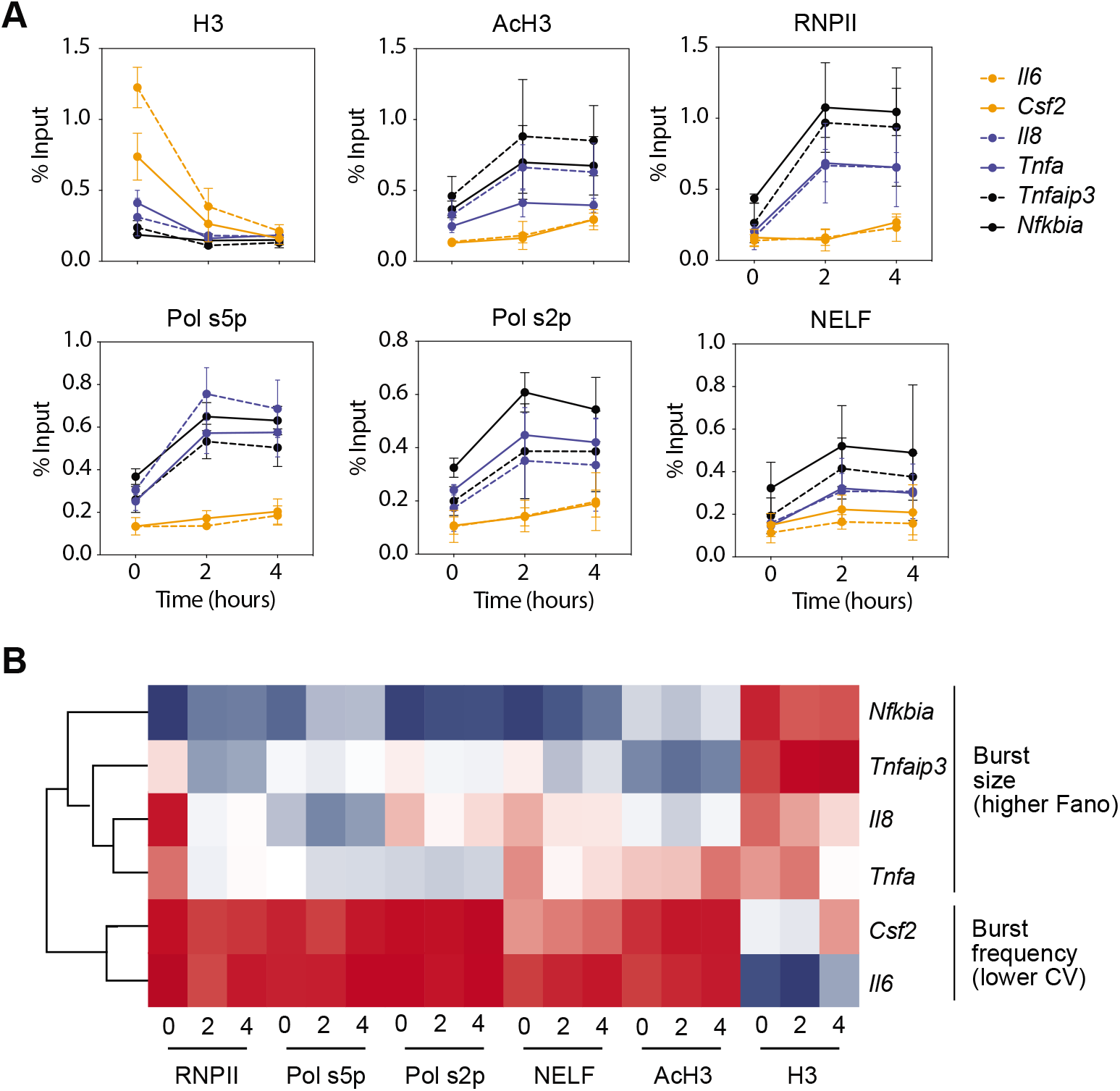
RNAPII pausing is associated with increases in transcriptional bursting upon TNF treatment. **A** Enrichment of histone H3, AcH3, total RNPII, ser5-p RNPII, ser2-p RNPII, and NELF-E in the basal state (0 hours) and after treatment (2 and 4 hours) with 20 ng/ml TNF quantified using ChIP and shown as % input (non-IP control). Data are presented as mean ± SD of three biological replicates. **B** Hierarchical clustering of ChIP data before and after TNF treatment separates promoters with TNF-mediated increases in burst frequency or burst size.

We also examined histone H3 acetylation at the target promoters by measuring total and acetylated H3. After TNF treatment, all six genes increased their AcH3 and decreased total H3 (Fig. 4A). *Nfkbia, Tnfaip3, Tnf*, and *Il8* increased AcH3 to higher levels than *Il6* and *Csf2*, while these two secondary cytokines decreased in total H3 to the same levels as the other targets at 2 and 4 hours after TNF treatment. Overall, there do not appear to be major differences in regulation of histone acetylation directly linked to the differences in bursting regulation, as all targets undergo similar changes in histone occupancy and regulation after TNF treatment. However, because chromatin remodeling is a molecular step that likely occurs before RNAPII regulation, basal differences in histone acetylation might underlie the differential changes we see in bursting.

Clustering our ChIP data, we find clear separation between *Il6* and *Csf2* and the more highly activated targets that show significant increases in burst size (Fig. 4B). Within the nonburst frequency increasing genes, *Nfkbia* separates from all other genes due to its increased accumulation of RNAPII, and the primary cytokines *Tnf* and *Il8* separate out from *Tnfaip3*. The clustering supports the idea that differences in molecular events occur at promoters of genes that have increased burst frequency (*Csf2*, *Il6*), burst size (*Tnfaip3*, *Tnf*, *Il8*), and highly activated genes that do not fit our mathematical model of bursting (*Nfkbia*).

### Small molecule inhibitors of histone acetylation and RNAPII pause release alter TNF-modulated transcriptional bursting

Our ChIP data suggested associations between basal H3 acetylation, RNAPII pausing and transcriptional bursting in response to TNF. To more directly test these associations, we perturbed these processes using small molecule inhibitors and measured effects on TNF-induced bursting for *Tnfaip3* and *Tnf*, which fall into the high and middle range of basal AcH3:H3 ratios among our targets (Fig. 1B).

To perturb histone acetylation, we pretreated Jurkat cells with the histone acetyltransferase (HAT) inhibitor A-485, a specific inhibitor of the HATs p300/CBP that are recruited by NF-κB (Fig. 5A) (Gerritsen *et al*, 1997; Lasko *et al*, 2017). We found that treatment with A-485 for 4 hours decreased AcH3 levels at the *Tnfaip3* and *Tnf* promoters but did not significantly affect total H3 levels as measured by ChIP-qPCR (Fig. 5B). A-485 pretreatment significantly decreased mean mRNA expression for both *Tnfaip3* and *Tnf* in response to 1 hour of TNF treatment (Fig. 5D). Pretreatment did not significantly alter CV or Fano factor for *Tnfaip3* or CV for *Tnf* after TNF treatment, but did reduce Fano for *Tnf* to basal levels (Fig. EV4A, B). Comparing single-cell mRNA distributions, the overall decrease in expression caused by A-485 was marked by an increase in *Tnf* non-expressing cells (Fig. EV4C), and a large reduction in the number of cells expressing much higher than the mean for both genes (Fig. 5D). These changes were more pronounced for *Tnf*, for which A-485 pretreatment completely eliminated the long-tailed distribution of cells expressing high numbers of mRNA, consistent with its impact on *Tnf* Fano factor.

**Figure 5.**
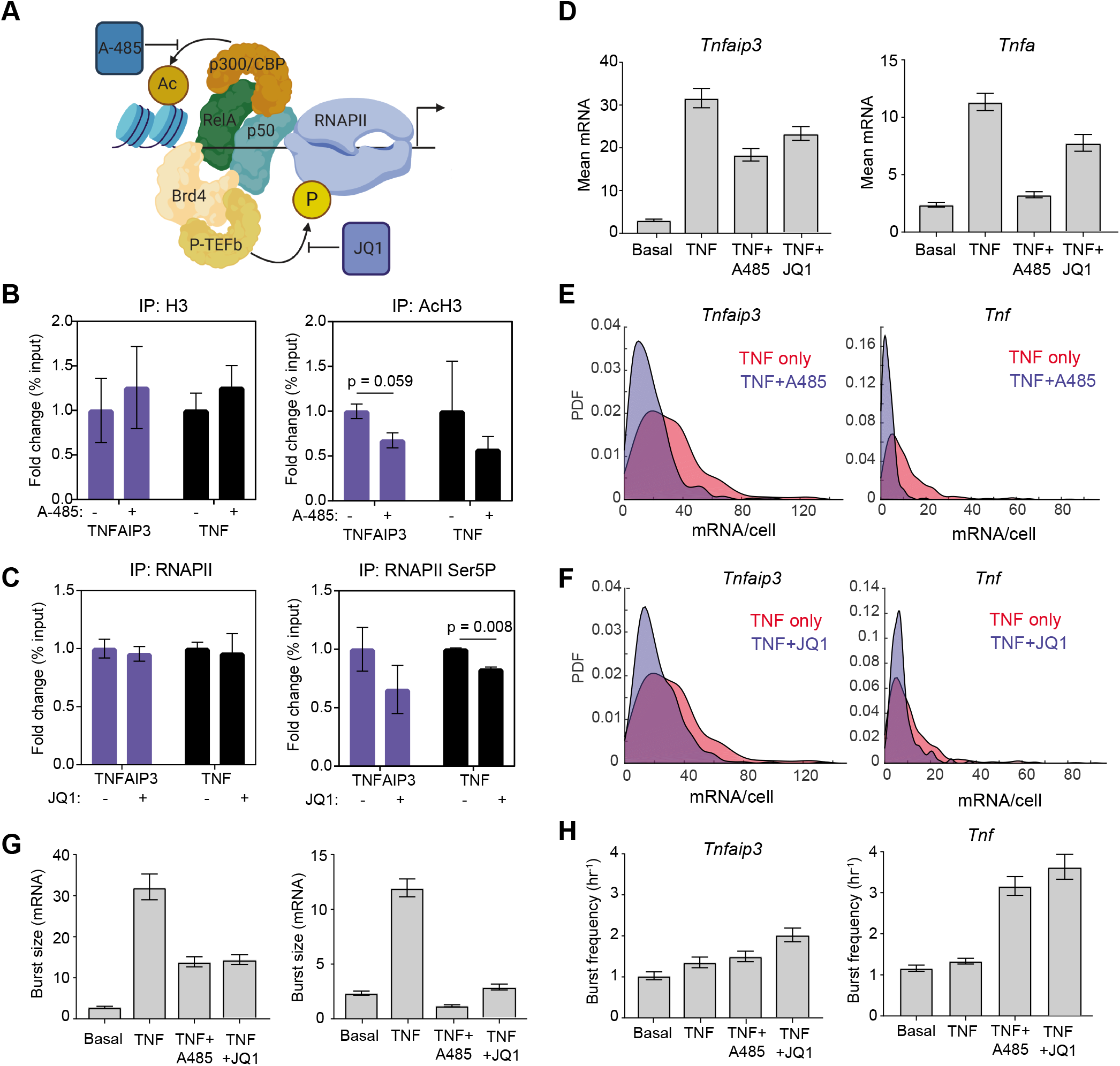
Small molecule inhibitors of H3 acetylation and RNAPII pause release alter TNF-mediated changes in transcriptional bursting. **A** Schematic of A-485 inhibition of the histone acetyl transferase p300/CBP, which is recruited by NF-κB and of JQ1 inhibition of BET bromodomains, which recruit the positive transcription elongation factor (P-TEFb). **B** Change in enrichment of histone H3 and AcH3 after treatment with 300 nM A-485 for 4 hours measured by ChIP-qPCR and shown as % input (non-IP control) normalized to the uninhibited control for each gene. Data are presented as mean ± SD of two biological replicates. p-values determined by t-test and reported if < 0.1. **C** Change in enrichment of total and Serine-5-phosphorylated RNAPII after 20ng/mL TNF treatment for 1 hour with 62.5nM JQ1 measured by ChIP-qPCR and shown as % input (non-IP control) normalized to the uninhibited control for each gene. Data are presented as mean ± SD of two biological replicates. p-values determined by t-test and reported if < 0.1. **C** Bar graphs of mean mRNA level in the basal state, or with and without A-485 pretreatment followed by 1-hour TNF treatment measured by smFISH for the indicated genes. Data are presented as mean ± bootstrapped 95% CIs. Samples with non-overlapping CIs are significant. **D** Bar graphs of mean mRNA in the basal state, or after TNF treatment for 1 hour with or without the indicated drug treatments measured by smFISH for the indicated genes. Data are presented as mean ± bootstrapped 95% CIs. Samples with non-overlapping CIs are significant. **E** Probability density of mRNA distributions measured by smFISH after 1-hour treatment with 20ng/mL TNF with (blue) or without (red) a 4-hour pretreatment with 300nM A-485. **F** Probability density of mRNA distributions measured by smFISH after 1-hour treatment with 20ng/mL TNF with (blue) or without (red) a 1-hour cotreatment with 62.5nM JQ1. **G** Burst size parameter fits from the two-state model in the basal state or after 1 hour 20 ng/mL TNF treatment with or without the indicated drug treatments. Data are presented as mean ± bootstrapped 95% CIs. **H** Burst frequency parameter fits from the two-state model in the basal state or after 1 hour 20 ng/mL TNF treatment with or without the indicated drug treatments. Data are presented as mean ± bootstrapped 95% CIs.

To perturb RNAPII pause regulation, we treated Jurkat cells with JQ1, an inhibitor of the BET family of bromodomain proteins, including BRD4, which recruits the positive transcription elongation factor b (p-TEFb) that stimulates pause release to NF-κB (Fig. 5A) (Filippakopoulos *et al*, 2010; Hargreaves *et al*, 2009; Huang *et al*, 2008). JQ1 has been found to decrease promoter-proximal RNAPII as well as alter the bursting of constitutively expressed genes (Bartman *et al*, 2019). We found that JQ1 decreased ser5-p RNAPII accumulation at the *Tnfaip3* and *Tnf* promoters as measured by ChIP-qPCR, but did not affect total RNAPII (Fig. 5C). JQ1 concomitantly decreased TNF-stimulated mean expression of both *Tnfaip3* and *Tnf* (Fig. 5D). JQ1 did not significantly alter CV or Fano factor for *Tnfaip3*, but significantly reduced both for *Tnf* (Fig. EV4A, B). When we compared the effect of JQ1 on single-cell mRNA distributions after 1-hour TNF treatment, we found that JQ1 decreased the number of cells in the high expressing tail of both *Tnfaip3* and *Tnf* without increasing the number of non-expressing cells for either gene (Fig. 5F and Fig. EV4C). This is in contrast to A-485, and is consistent with our observation that only ser5-p RNAPII, but not total RNAPII, decreased at the target promoters. Overall, these inhibitors decrease TNF-stimulated transcription while differentially affecting mRNA distributions.

Fitting mRNA distributions for TNF treatment following pretreatment with A-485 to the theoretical pdf of the two-state model, we found that, in response to TNF treatment, A-485 decreased burst size for *Tnfaip3* while not affecting burst frequency. For *Tnf*, A485 pretreatment prior to TNF stimulation caused an increase in burst frequency without any change in burst size compared to the basal state (Fig. 5G, H). Overall, A-485 pretreatment reshaped TNF-induced changes in transcriptional bursting such that the transcriptional bursting response of *Tnfaip3* resembles *Tnf* and *Il8* (no change in burst frequency combined with a smaller increase in burst size) and the response of *Tnf* resembles *Il6* and *Csf2* (no change in burst size combined with an increased burst frequency). JQ1 also reduced TNF-induced burst size increases for both *Tnfaip3* and *Tnf*, and this was accompanied by an increase in burst frequency for both genes, although this change was more pronounced for *Tnf* (Fig. 5G, H). These observations support the hypothesis that basal histone H3 acetylation levels at NF-κB target promoters affect how TNF treatment alters transcriptional bursting, and that TNF-stimulated accumulation of paused RNAPII at its target promoters is linked to an increase in transcriptional burst size.

### Mathematical modeling shows how TNF positive feedback amplifies skewed distributions produced by transcriptional bursting to create more heterogeneous cell populations

TNF modulates transcriptional burst size at some promoters and burst frequency at others, producing more or less skewed mRNA distributions across a cell population, respectively. In addition, TNF positively regulates its own production, and our results show that it does this by increasing transcriptional burst size. We and others have shown that TNF exhibits significant cell-to-cell variability in response to NF-κB stimulation by TNF and also EPS (Adamson *et al*, 2016; Bagnall *et al*, 2018; Sung *et al*, 2014; Xue *et al*, 2015). Therefore, we sought to explore how modulation of burst size combined with positive feedback contributes to this observed heterogeneity.

We built a mathematical model of a two-state promoter responding to an initial TNF stimulus. As TNF can stimulate its own transcription, we included positive feedback from the newly produced TNF on the cell that produced it (simulating autocrine signaling) to explore its effects on cell-to-cell heterogeneity (Fig. 6A). We modeled the addition of exogenous TNF as a time-dependent change in k_t_, the mRNA production rate, in order to match the change in burst size inferred from our smFISH distributions. We fit this k_t_ function empirically to reproduce our averaged experimental mRNA data (Fig. EV5A). We explored a range of TNF positive feedback parameters and their effect on the level and timing of maximum *Tnf* (Fig. EV5B-E). We chose values that qualitatively reproduced the dynamic changes in cell-population averages and distributions observed in our population level RT-qPCR measurements of *Tnf* transcription (Fig. 6B and Fig. 1C). We confirmed that including positive feedback increased and sustained the peak of *Tnf* mRNA at 1 hour and TNF protein production at 8 hours (Fig. 6B and C).

**Figure 6.**
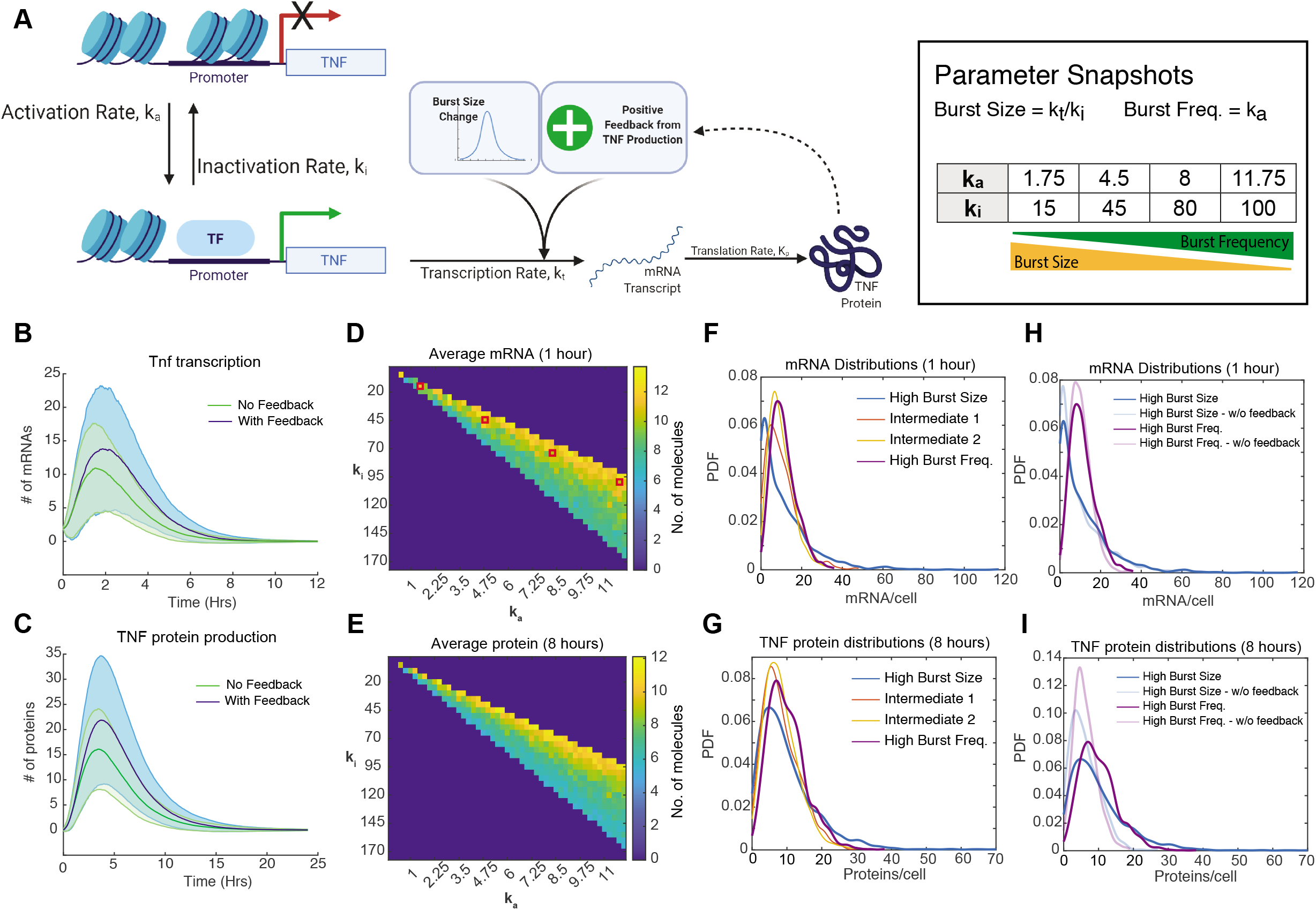
Mathematical modeling demonstrates the functional consequences of increasing burst size versus burst frequency after TNF stimulation. **A** Schematic of the two-state model of transcription coupled to translation of a protein (TNF) that positively feeds back on its own transcription rate. **B, C** Model simulation of cell-population averages of *Tnf* mRNA (B) and TNF protein (C) versus time with and without positive feedback. Data are presented as mean (dark line) and SD (shaded region) of 1,000 simulated cells. **D, E**: Cell-population averages from stochastic simulations with positive feedback of *Tnf* mRNA at 1 hour (D) and TNF protein at 8 hours (D) after TNF treatment across a parameter space with increasing burst frequency (k_a_) and decreasing burst size (k_i_) chosen to produce levels of *Tnf* mRNA similar to experimental observations. **F, G**: Probability density function plots of simulated single-cell mRNA (F) and protein (G) numbers for the indicated bursting parameters in response to exogenous TNF treatment. **H, I**: Probability density function plots of simulated single-cell mRNA (H) and protein (I) numbers with and without the inclusion of positive feedback for either an increase in only burst size or only burst frequency in response to exogenous TNF treatment.

Our goal was to use this model to compare how positive feedback affected cell-to-cell heterogeneity of TNF production when amplifying TNF-stimulated transcriptional increases of similar means but with different noise. To do this, we performed a parameter scan in the absence of positive feedback, in which we increased burst frequency (i.e., the activation rate k_a_) and simultaneously decreased burst size (i.e., by increasing the inactivation rate k_i_). By increasing burst frequency while simultaneously decreasing burst size, we were able to identify a region in which mean expression remains relatively constant but noise varies due to differences in burst behavior (Fig. 6D and E).

To analyze how cell-to-cell heterogeneity of TNF production changed with different bursting behaviors, we chose four parameter sets. These included one with a large change in burst size and no change in burst frequency in response to TNF that resembles our experimental data, one with a large increase in burst frequency and no change in burst size to explore the opposite extreme, and two with intermediate changes in both. As seen in our experimental data, increasing burst size created more long-tailed single-cell mRNA distributions, though the effect was somewhat lessened at the protein level (Fig. 6F and G). Quantifying the mean, CV, Fano factor, and skew of these distributions, we observed that as transcription activity shifted from burst size to burst frequency changes, mean remained constant, while CV, Fano, and skew all decreased (Fig. EV5F).

To examine the effect of TNF positive feedback on its own production, we compared simulations of the parameters sets that increased either burst size or burst frequency with and without positive feedback. Including positive feedback produced relatively small increases in mean mRNA levels, but these differences were amplified at the protein level (Fig. EV5F). Positive feedback also increased the length of the distribution tails, and importantly it did this more for the burst size-increasing condition than for the burst frequency-increasing condition (Fig. 6H-I). In other words, a small population of high TNF-producing cells was more pronounced when transcription was increased via burst size vs. burst frequency. The effect of positive feedback on increasing cell-to-cell heterogeneity was evident in the large increase in Fano factor that was greatest for the burst size-increasing condition (Fig. EV5F). In contrast, including feedback had very little effect on CV (Fig. EV5F). Overall, our modeling indicates that positive feedback by TNF coupled with transcriptional increases driven by burst size modulation leads to highly skewed distributions of protein across cells. We speculate that these mechanisms contribute to small subpopulations of cells with high functionality, such as high TNF-producing cells that have been observed in response to activation of the NF-κB-mediated inflammatory response in other studies.

## Discussion

Transcriptional bursting is an important process affecting many biological processes, but it has not been extensively studied for endogenous NF-κB targets, including cytokines that are vital to the inflammatory response. Here, we explored changes in transcriptional bursting in response to the inflammatory cytokine TNF in T cells. We found that TNF can modulate either burst frequency or burst size depending on basal histone acetylation and regulation of RNAPII pausing. Using small molecule inhibitors, we confirmed that altering basal histone acetylation or RNAPII pausing after TNF stimulation altered bursting behavior for *Tnfaip3* and *Tnf*. Finally, we used mathematical modeling to show that increasing burst size in response to TNF can lead to more skewed single-cell distributions as compared to increasing burst frequency and this is substantially amplified with TNF positive feedback.

We found that TNF primarily increased transcription by increasing burst size, which resulted in highly skewed, long-tailed mRNA distributions that are generally marked by large increases in Fano factor. In contrast, transcription factor-mediated increases in burst frequency result in less skewed distributions with lower cell-to-cell heterogeneity. Increases in burst frequency in response to transcription factor stimulation have been more commonly observed than increases in burst size (Chen *et al*, 2019; Friedrich *et al*, 2019; Li *et al*, 2018). Notably, most of these examples analyzed cellular processes for which it is important that most or all cells in a population respond to a stimulus with similar levels of gene expression such as the DNA damage response (Friedrich *et al*, 2019) or the circadian response to light (Li *et al*, 2018). In contrast, for processes where highly skewed single-cell responses are beneficial, as appears to be the case for inflammatory signaling, stimulus-induced burst size increases may be more common. Long-tailed distributions with a few outliers far above the population mean have been shown to be important for regulating inflammatory signaling at the levels of single-cell transcription (Shalek *et al*, 2014) and cytokine secretion (Xu *et al*, 2015). Burst size regulation was also observed in response to Notch signaling, which is active in both embryonic development and maintenance of the germline stem cell niche (Falo-Sanjuan *et al*, 2019; Lee *et al*, 2019). Creating skewed distributions in transcription between cells that must follow different trajectories such as proliferation versus differentiation might help ensure that cells do not easily cross over to the other behavior.

We found that accumulation of promoter-proximal paused RNAPII in response to TNF was linked to increased burst size. RNAPII promoter-proximal pausing occurs throughout the mammalian genome, especially at signal-responsive promoters (Adelman and Lis, 2012). Paused RNAPII primes a promoter to rapidly respond to an elongation signal, bypassing the need to recruit a new RNAPII subunit. In our study, accumulation of paused RNAPII occurred at genes with higher basal histone acetylation. We have previously found that both basal histone acetylation and RNAPII pause regulation after TNF treatment are linked in the same ways to transcriptional bursting from the HIV LTR promoter (Wong *et al*, 2018). The accumulation of RNAPII may also prime promoters to respond more strongly to additional pro-inflammatory stimuli, either cytokines produced in response to the first stimulus or another external source.

To model transcriptional bursting, we used a simple two-state model in which a promoter can occupy either an active ‘on’ state or inactive ‘off’ state, which fit our data from Jurkat T cells well. Including a third refractory promoter state that an active promoter transitions to before transitioning back to the inactive state has produced better fits in some systems, including bursting of the HIV LTR promoter in HeLa cells (Li *et al*, 2018; Suter *et al*, 2011; Zambrano *et al*, 2020). However, the extremely low transition rates we observed from the inactive to the active state (burst frequencies on the order of 1 per hour) might be slower than (and thus mask) transitions from a refractory to the inactive state. Identification of endogenous NF-κB targets with much higher burst frequencies, possibly in other cell types, would help test if a more complex model configuration more accurately explains transcription of NF-κB targets.

HIV encodes its own positive feedback mediator, the protein Tat, which leads to strong positive feedback and full HIV activation in long-tailed distributions that stem from burst size increases (Wong *et al*, 2018). The inflammatory cytokine TNF also positively regulates its own expression either in the producing cell or in neighboring cells via autocrine or paracrine signaling. We used mathematical modeling to explore how the shape of the single-cell *Tnf* mRNA distribution might be related to biological function. We found that increasing burst size rather than burst frequency is required to create a highly skewed single-cell transcript distribution, and this is amplified by positive feedback. Positive feedback had a much smaller effect on distributions from a burst frequency increase. We only accounted for autocrine signaling in our model, and did not consider how the TNF produced by one cell might affect neighboring cells. However, paracrine signaling plays a major role in regulating immune signaling, so signals from the cells in the high expressing tail of a distribution could have a significant role in propagating a pro-inflammatory response *in vivo*. More work will need to be done to explore the role of transcriptional bursting in inflammatory gene expression in other immune cell types and for other stimuli besides the cytokine TNF.

## Methods

### Reagents and tools table

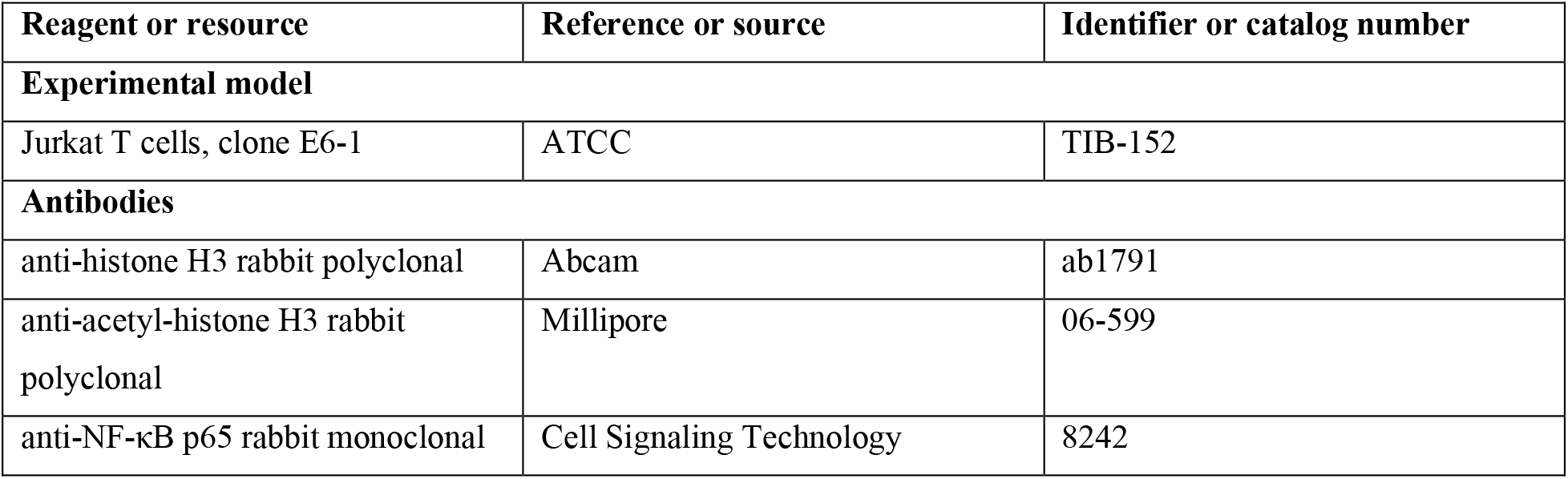
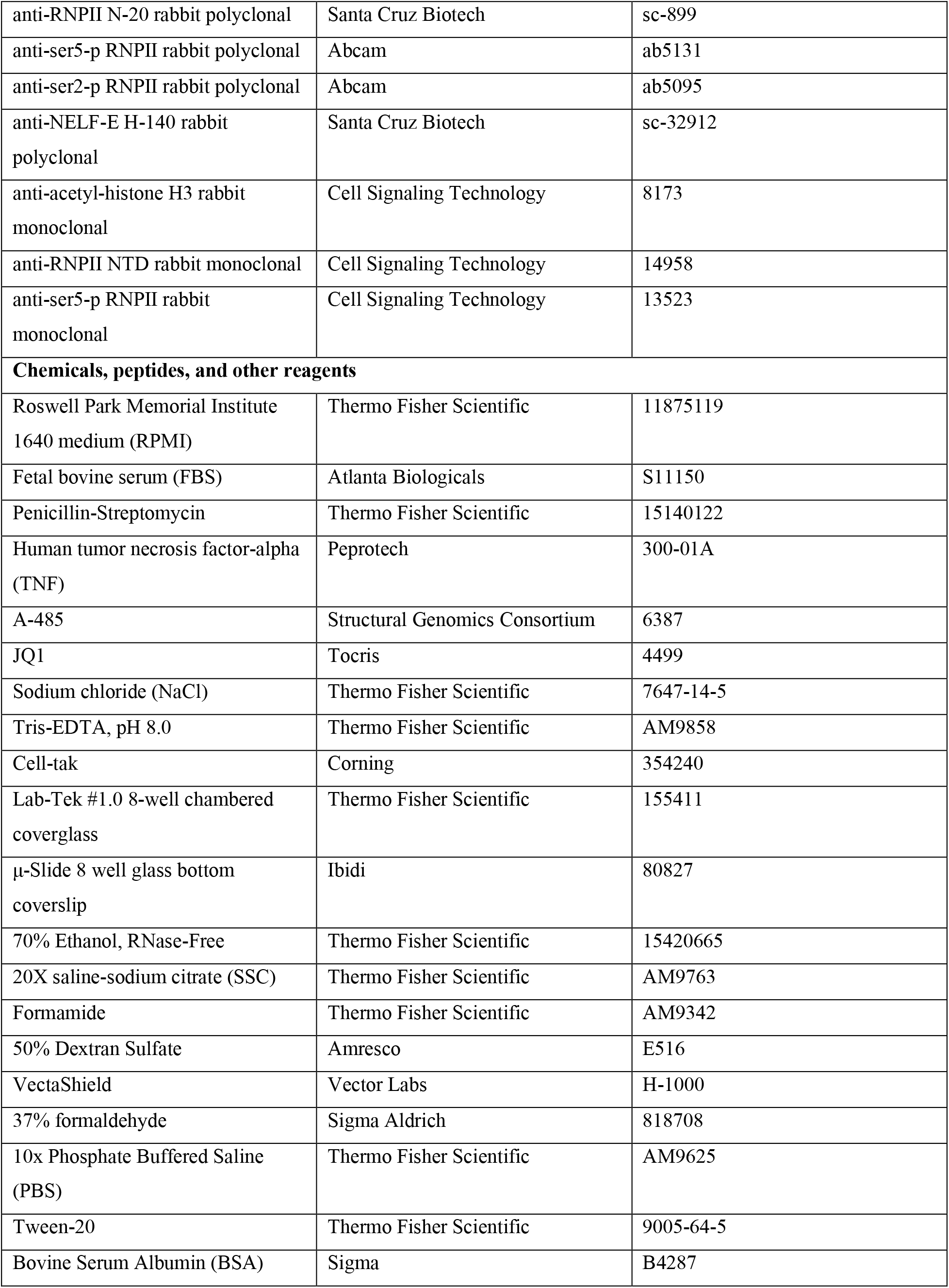
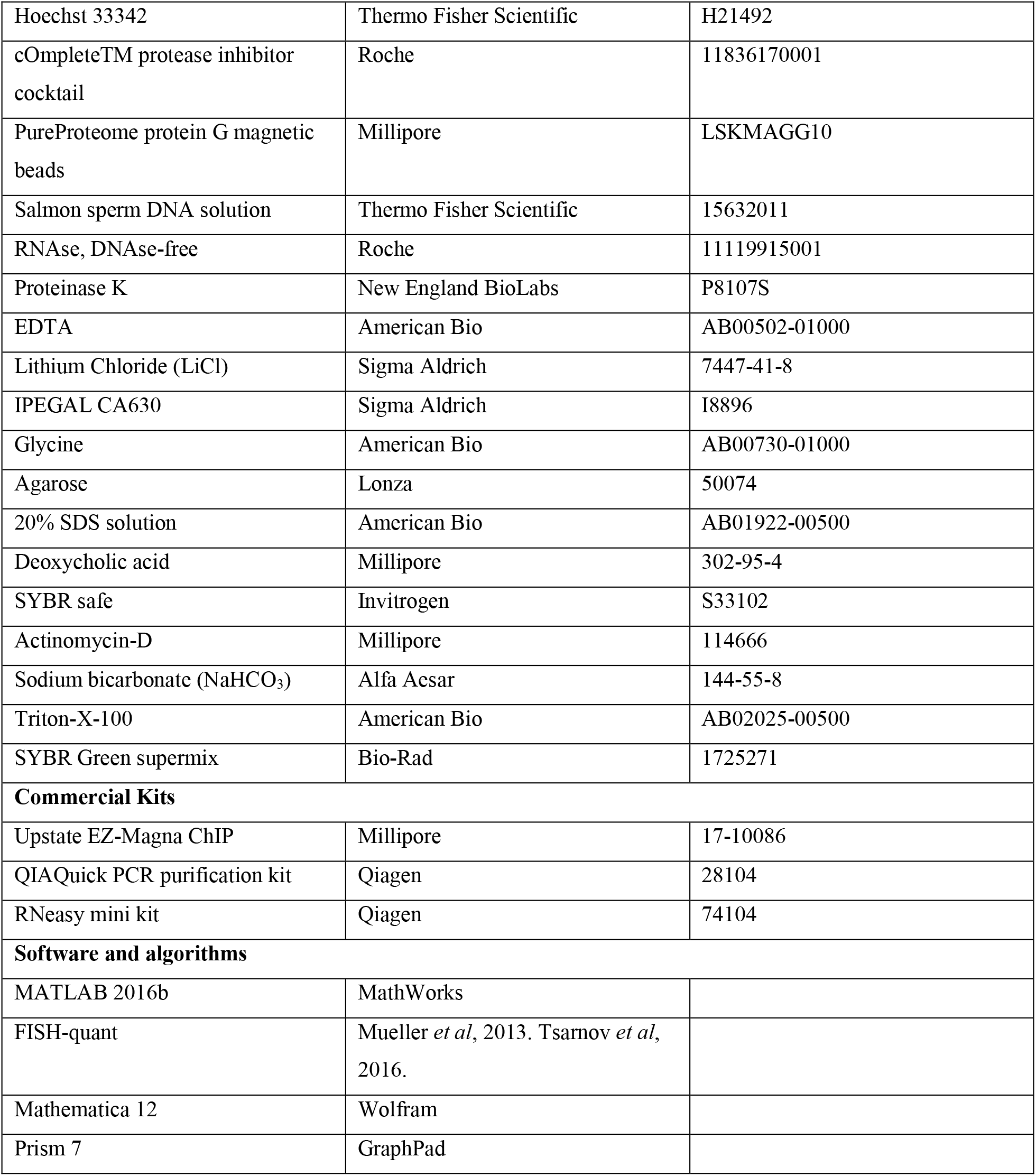

### Cell culture and pharmacological treatments

Jurkat T cell clone E6-1 were obtained from ATCC. Jurkat cells were cultured in Roswell Park Memorial Institute 1640 (RPMI) medium (Thermo Fisher Scientific). All media was supplemented with 10% fetal bovine serum (Atlanta Biologicals), 100 U/mL penicillin, and 100 μg/mL streptomycin (Thermo Fisher Scientific). Cells were maintained in 5% CO_2_ at 37°C and were never cultured beyond passage 20. Cells were grown to at least 500,000 cells/mL before treatment with 20 ng/mL recombinant human tumor necrosis factor α (TNF) (Peprotech), 300 nM A-485 (Structural Genomics Consortium), or 62.5 nM JQ1 (Tocris). To calculate mRNA decay rates, Jurkat cells were stimulated with TNF for 1 hour followed by 10 μg/mL actinomycin-D treatment for varying times.

### Chromatin immunoprecipitation

Chromatin immunoprecipitation was performed using the Upstate EZ-Magna ChIP kit (Millipore). Briefly, 5 million cells per condition were fixed in 1% formaldehyde (Sigma) for 10 minutes, after which excess formaldehyde was quenched with 10X glycine at room temperature. Cells were washed three times with ice cold PBS and then lysed in 300 μL of 1% SDS lysis buffer with protease inhibitor cocktail (Roche). Lysates were sonicated with a Diagenode Bioruptor Plus with the following settings: 30 minutes of 30 seconds ON/30 seconds OFF at high power in a 4°C water bath. Sheared DNA was run on a 1% agarose gel (Lonza) to verify that sheared DNA was between 100-1000 bp. Samples were pre-cleared with PureProteome Protein G magnetic beads (Millipore) at 4°C and 5% of each sample was aliquoted as a percent input control. Samples were incubated with antibody at manufacturers’ recommended concentrations overnight at 4°C. PureProteome beads were added and incubated for one hour at 4°C. Beads were washed once each with low salt, high salt, and LiCl immune complex wash buffers, then washed twice with TE buffer, and then eluted with elution buffer at room temperature. Crosslinks were reverse by incubating samples with NaCl overnight at 65°C. DNA was purified using the QIAQuick PCR Cleanup kit (Qiagen). DNA was quantified using quantitative PCR using SsoAdvanced Universal SYBR Green Supermix on a CFX Connect Real-Time System (BioRad). qPCR was run in triplicate and melt curves were run to confirm product specificity.

### RT-qPCR

Total RNA was purified with the RNeasy Mini kit (Qiagen), including an on-column DNase treatment. cDNA was synthesized using SuperScript III reverse transcriptase (Thermo Fisher Scientific) and dT oligo primer. cDNA was diluted in nuclease-free water and quantified using SsoAdvanced Universal SYBR Green Supermix on a CFX Connect Real-Time System (BioRad) with the following amplification scheme: 95°C denaturation for 90 seconds followed by 40 cycles of 95°C for 15 seconds, 60°C annealing for 10 seconds, and 72°C elongation for 45 seconds with a fluorescence read at the end of each elongation step. This was followed by a 60°C to 90°C melt-curve analysis with 0.5°C increments to confirm product specificity. All samples were normalized to the house-keeping gene *Gapdh*.

### smFISH probe design, hybridization, and imaging

The probe sets targeting *Nfkbia*, *Tnfaip3*, and *Il8* (Lee *et al*, 2014) and *Tnf* (Bushkin *et al*, 2015) were previously described. The probe sets targeting *Il6* and *Csf2* were designed using the Stellaris^®^ RNA FISH Probe Designer (Biosearch Technologies, Inc., Petaluma, CA) available online (www.biosearchtech.com). All mRNAs were hybridized with Stellaris RNA FISH Probes labeled with Quasar 670 (Biosearch Technologies, Inc.) following the manufacturer’s instructions. Briefly, Jurkat cells were treated under indicated conditions and then plated onto Cell-Tak (Corning) coated Lab-Tek #1.0 8-well chambered coverglass (Thermo Fisher Scientific) or μ-Slide 8 well glass bottom coverslip (Ibidi). Cells were fixed in 3.7% formaldehyde (Thermo Fisher Scientific) for 10 minutes and then permeabilized overnight in 70% ethanol (Fisher Scientific). Cells were hybridized for 12 hours overnight with the following probe set specific conditions: 250 nM probe for *Tnf/Il6/Csf2* in 2X SSC (Thermo Fisher Scientific) with 10% formamide (Thermo Fisher Scientific) and 100 mg/mL dextran sulfate (Amresco) at 37°C, 50 nM probe for *Tnfaip3* in 2X SSC with 10% formamide and 80 mg/mL dextran sulfate at 37°C, 250 nM probe for *Nfkbia* in 2X SSC with 12% formamide and 100 mg/mL dextran sulfate at 37°C, and 250 nM probe for *Il8* in 2X SSC with 10% formamide and 100 mg/mL dextran sulfate at 25°C. After hybridization, cells were washed twice with 2X SSC and 10% formamide, counterstained with 100 ng/mL Hoechst 33342 (Thermo Fisher Scientific) for 15 minutes, and immersed in VectaShield mounting media (Vector Labs). Cells hybridized with *Nfkbia*, *Tnfaip3*, *Tnf*, and *Il6* probes were imaged on an Axio Observer Zi inverted microscope (Zeiss) with an Orca Flash 4.0 V2 digital CMOS camera (Hamamatsu) and a 100x APO oil objective (NA 1.4, Zeiss). Cells hybridized with *Il8* and *Csf2* probes were imaged on an LSM 510 spinning disk confocal microscope (Zeiss) with a C9100-13 camera (Hamamatsu) a 100x TIRF APO oil objective (1.49 NA, Nikon)

### smFISH image analysis

We quantified mRNAs in individual cells using FISH-Quant in MATLAB R2016B (Mathworks Inc.) (Mueller *et al*, 2013, Tsanov *et al*, 2016). Cells were manually identified and outlined, with overlapping cells, cells partly in the field of view, and multinucleated cells excluded from analysis. Images for all probe sets were filtered using the 2x Gaussian filtering method in FISHQuant. Thresholds for mRNA spot detection were determined for each image set by testing the FISH-Quant software on images of high and low expressing cells (TNF treated and untreated) and comparing to values that were visually derived. The remaining images were then processed in batch. Pre-detection thresholds and detections settings were set for each experimental condition.

### Fitting the two-state model

Maximum-likelihood estimation (MLE) was used to select burst frequency (k_a_) and burst size (b = k_t_/k_i_) parameters that best fit the measured mRNA distributions to the full analytical solution to the two-state stochastic gene expression model (Peccoud and Ycart, 1995). MLE was performed as numerical minimization over the negative log-likelihood function defined over the probability density function (pdf) given the observed experimentally determined RNA distributions for each condition using the method of moments. As previously reported, mRNA distributions are not sufficient to independently determine the promoter inactivation rate k_i_ and the transcription rate k_t_. Using a previously described method (Rat *et al*., 2006. Skupskey *et al*., 2010), we held the transcription rate k_t_ constant across all conditions and reported b. Sensitivity analysis of the k_t_ value for each gene suggested that our results are largely independent of the k_t_ value chosen for each gene (Fig. EV2B). MLE was implemented using custom code in Mathematica 8 (Wolfram Inc.) as previously described (Dey *et al*., 2015).

### Statistical analysis

All smFISH experiments included a sufficient number of cells to characterize the transcript distributions (n > 100 cells) and results were confirmed with independent biological replicates. The 95% confidence intervals on all descriptive statistics of RNA distributions were estimated from the 2.5% and 97.5% quantiles of bootstrapped copy number counts per cell as previously described (Dey *et al*., 2015). 95% confidence intervals on fit burst frequency and size parameters were estimated from the log-likelihood function assuming asymptotic normality of the estimates and using 1.92 log-likelihood ratio units as previously described (Dey *et al*., 2015). The difference between two quantities was inferred to be significant (p < 0.05) if the 95% CI’s were not overlapping (Schenker and Gentleman 2001). All regression and correlation analysis was performed in Prism (Graphpad).

### Mathematical Model Development

We modified a previously published mathematical model of a two-state promoter with positive feedback (Wong *et al*., 2018). Briefly, we modeled transcription as a promoter that transitions from an ‘OFF’ state to an ‘ON’ state, and vice versa, with rate constants, k_a_ and k_i_ respectively. In the ‘ON’ state, mRNA is produced at the rate k_m_ and degraded at a rate of g_m_. This rate was modulated by time via a fitted burst size curve (see below). The mRNA produces TNF protein at the rate k_p_, is exported out of the cell at a rate k_ex_, and degraded at a rate of g_p_. To model TNF positive autoregulation, a feedback loop was introduced into the model to increase the rate of mRNA production, k_m_, in respone to increasing exogenous TNF protein. The reactions governing this model and the rate constants are described in Table 1.

**Table 1:** Model reactions and rate constants

| Reaction | Parameter (units) | Values | Source |
| --- | --- | --- | --- |
| *Promotor On* → *Promotor Off* | $k_i(hr^{-1})$ | 15 | Range inferred from experimental distributions |
| *Promotor Off* → *Promotor On* | $k_a(hr^{-1})$ | 1.3 | Range inferred from experimental distributions |
| *Promotor On* * (*fm*) → *mRNA* | $k_t(hr^{-1})$ | $k_i * b$ | Calculated from fitted burst size equation ($b = k_m/k_i$) |
| $fm = 1 + \frac{A * [TNF(outside)]}{K + [TNF(outside)]}$ | $K, A$ | 500, 25 | Parameter scan |
| *mRNA* → *TNF (inside)* | $k_p(hr^{-1})$ | 1 | Estimated from (Caldwell *et al.*, 2014) |
| *TNF (inside)* → *TNF(outside)* | *export* ($hr^{-1}$) | 18 | (Paszek, *et al.*, 2010) with assumptions from (Lee, *et al.*, 2014) |
| *TNF* $\rightarrow \emptyset$ | $g_p(hr^{-1})$ | 0.36 | Estimated from mRNA degradation (1/3 of $g_m$) |
| *mRNA* $\rightarrow \emptyset$ | $g_m(hr^{-1})$ | 1.09 | Experimental derivation |

This model is represented by the following system of ordinary differential equations:

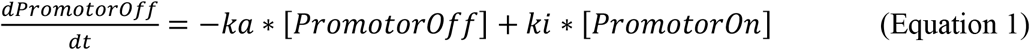

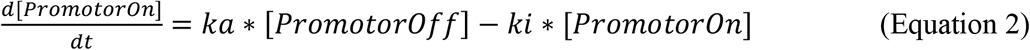

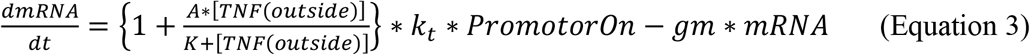

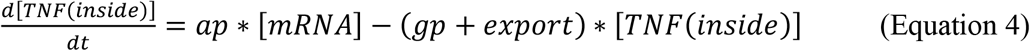

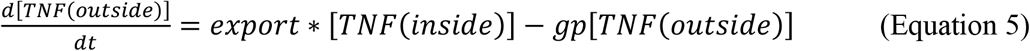

To stochastically simulate TNF protein and mRNA transcript production over time, we used a network free stochastic simulator (NFSIM) (Sneddon et al., 2011), and then analyzed and plotted outputs using MATLAB R2019B (MathWorks, Inc.).

### Steady-State Analysis

To understand how the system behaves under basal conditions, we assumed equilibrium for the above equations. First, we examined promotor dynamics, and solved for steady-state. By solving EQ 1 and 2 at steady-state, and setting

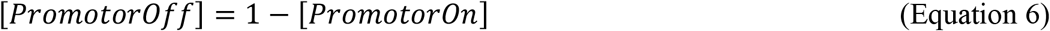

we derive the following:

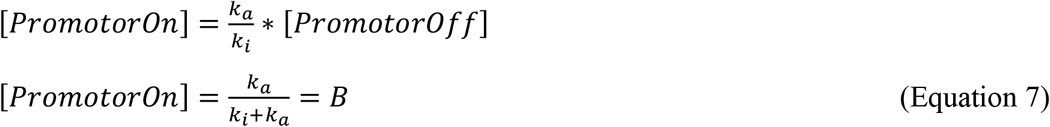

We can then examine TNF concentrations inside and outside the cell (EQ 4 and 5), by deriving the following:

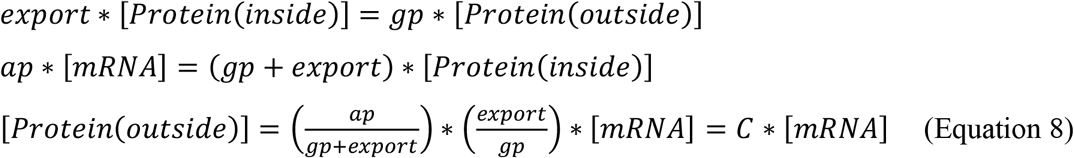

Finally, we use EQ3, EQ7, and EQ8 to solve for mRNA values under steady-state conditions, following:

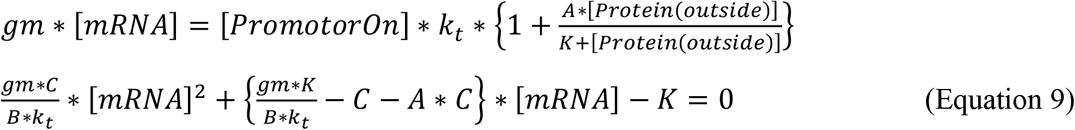

Using this equation, we explored how the parameter space affects mRNA concentration before TNF treatment. By varying feedback parameters–the amplification rate, and the K half-max–we recreated regions that matched basal *Tnf* mRNA conditions (Fig. EV5E). To understand how the unbounded feedback parameters influenced the model upon TNF treatment, we ran large 2-D parameter scans altering feedback parameters. Two parameters were chosen that qualitatively reproduced the time course of TNF-activation experiments under deterministic simulation of the model (Fig. EV5B-D) and replicated steady-state values of basal mRNA (Fig. EV5E).

To explore how changes in burst size and burst frequency influenced phenotypic outcomes, we stochastically simulated 1,000 cells using NFSIM, altering the parameters k_i_ and k_a_. Four representative parameter combinations were chosen, each with an average of 10 mRNA transcripts per cell at 1 hour (Fig. S1A-B). To depict phenotypic differences of these cells, the amount of mRNA transcripts and proteins were plotted as normal kernel density functions at 1 hour and 8 hours, respectively.

### Burst Size Phenomenological Equation

Addition of TNF increases experimental burst size in a time-dependent curve. As TNF is degraded or used, the burst size begins to decrease. We fit an exponential curve to our experimental values, weighting the fit according to variance of the data (Figure EV5A). This phenomenological equation reflects TNF’s mechanistic activation of NF-κB, and further promotion of transcription.

### Data availability

Quantitative smFISH measurements presented in the main figures are provided as figure source data. Modeling computer scripts are available at https://github.com/elisebullock/tnftwostate

## Acknowledgements

We thank Dr. Nadya Dimitrova for advice on chromatin immunoprecipitation, and Dr. Valerie Horsley and Dr. Yannick Jacob for use of lab equipment. We thank Dr. Sanjay Tyagi for sharing TNF smFISH probe sequences. We thank the Structural Genomics Consortium for providing the small molecule A-485. This work was funded by the National Science Foundation (CBET-1454301 to K.M.-J.), and the NIH (1R01-GM123011 to K.M-J.). V.L.B. was supported by NIH predoctoral training grants in virology (5T32AI055403-12 and 5T32AI055403-13). V.C.W. was supported by NIH predoctoral training grants in genetics (2T32GM007499-36, 5T32GM007499-34, and 5T32GM007499-35). M.E.B. was supported by NIH grant 1T32EB019941.

## Author contributions

V.L.B., V.C.W., S.G., and K.M-J. conceived the study and designed the experiments. V.L.B. and V.C.W. performed experiments. M.E.B. performed mathematical and stochastic modeling.

V.L.B., V.C.W., M.E.B., and K.M-J. analyzed data. V.L.B., M.E.B., and K.M-J. prepared figures and wrote the manuscript. All authors edited the manuscript. K.M-J. acquired funding and supervised the research.

---

**Figure EV1.**
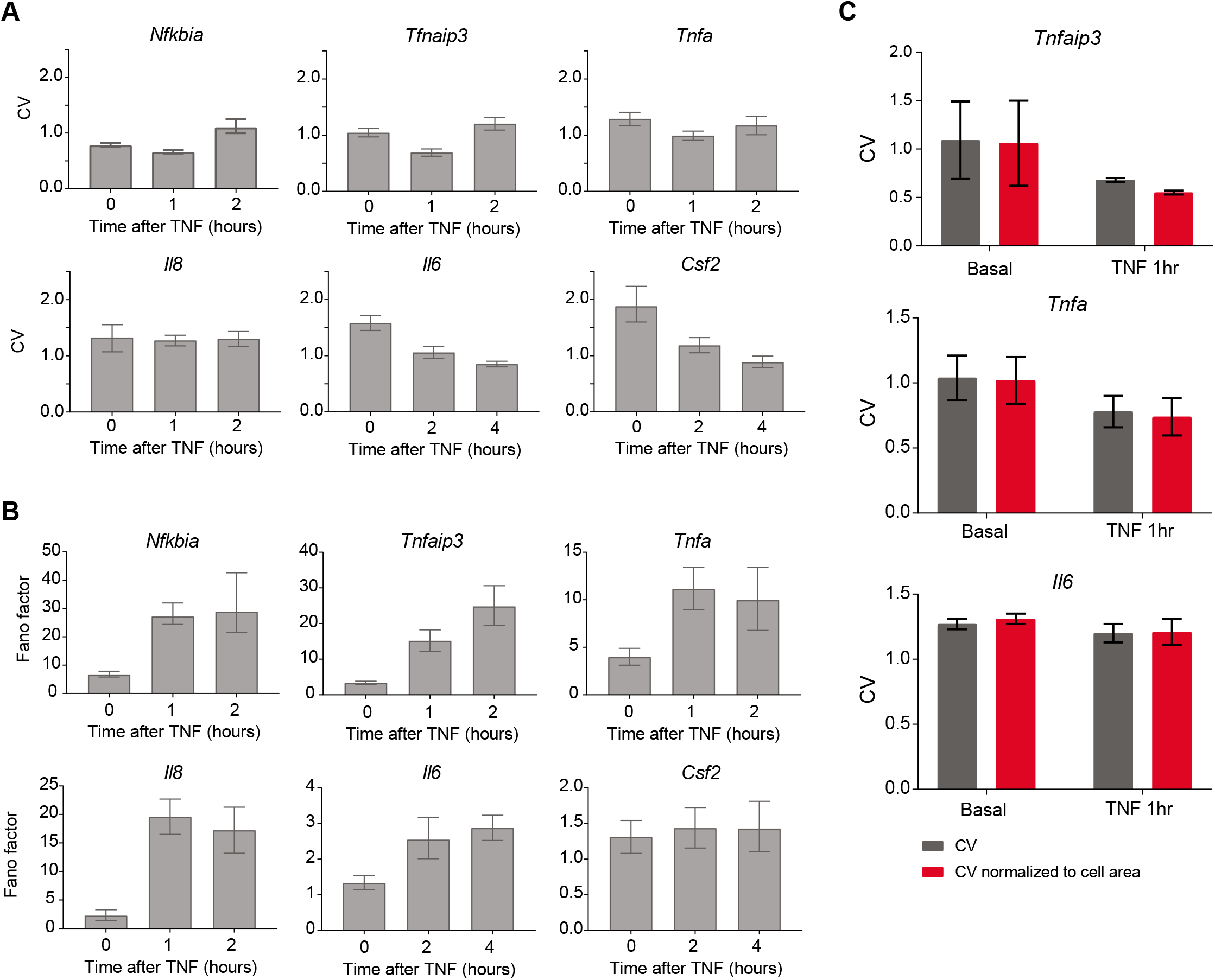
TNF treatment induces differential changes in noise across target genes. **A, B** Bar graphs of coefficient of variation (CV) (A) and Fano factor (B) of TNF-stimulated smFISH distributions presented in Fig. 2B for the indicated genes at the indicated time points. Data are presented as mean and bootstrapped 95% confidence intervals (CIs). **C** Coefficient of variation from mRNA distributions calculated from raw data (red) or mRNA counts normalized to cellular area (red) are shown. Data are presented as mean +/-standard deviation (SD) of CV’s from two replicate smFISH experiments for each gene and condition.

**Figure EV2 (related to Fig. 3).**
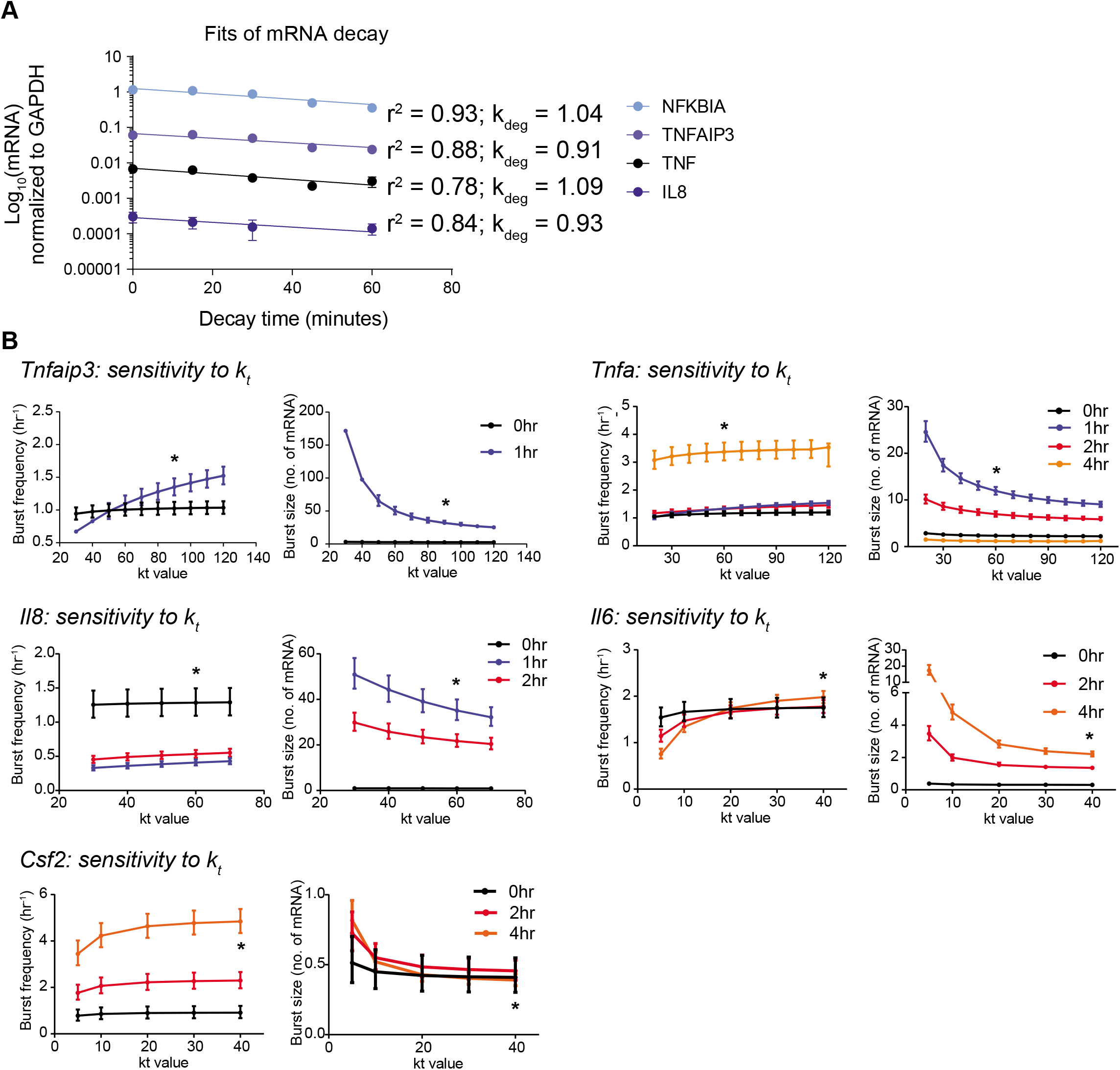
Validation of parameters used to fit burst size and burst frequency values from the two-state model. **A** Jurkat T cells were stimulated with 20 ng/mL TNF for 1 hour followed by treatment with 10 μg/mL Actinomy-cin-D for 0, 15, 30, 45, or 60 minutes and mRNA levels for the indicated target genes were measured by RT-qPCR. Exponential mRNA decay rates were fit using non-linear regression. Data are presented as mean +/- SD of three biological replicates. **B** Sensitivity analysis of how fitted burst frequency (left) and burst size (right) values vary with the value of kt used in the model. Data are presented as mean and bootstrapped 95% CIs. The k_t_ value used is indicated with an asterisk.

**Figure EV3 (related to Fig. 3).**
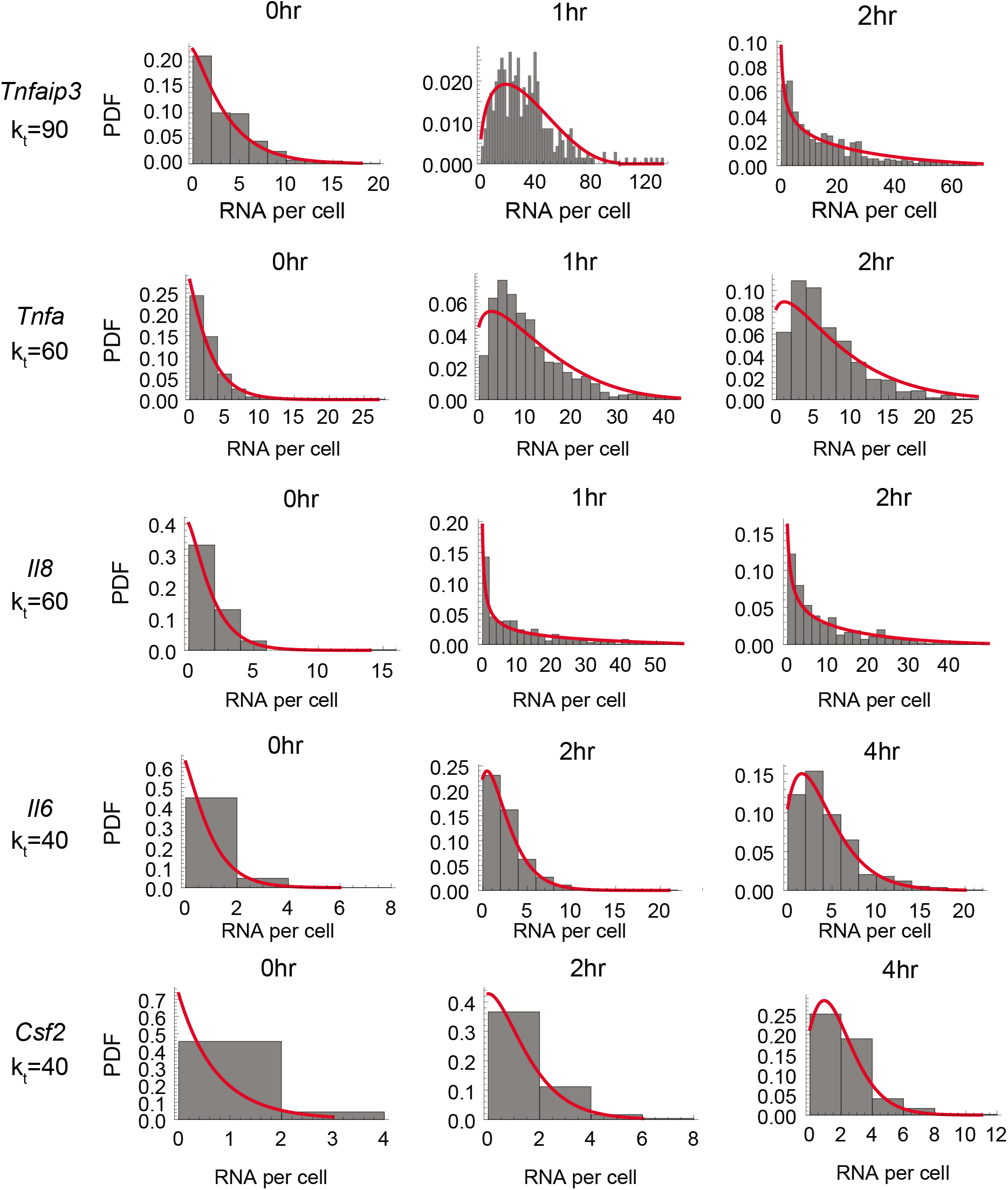
Probability density functions for best-fit two-state promoter model parameters. Histograms (grey) of mRNA distributions for the indicated genes measured by smFISH in the basal state and after 20 ng/mL TNF treatment. Red curves show the best fit analytical probability density function of a stochastic two-state promoter model found by maximum likelihood estimation.

**Figure EV4 (related to Fig. 5).**
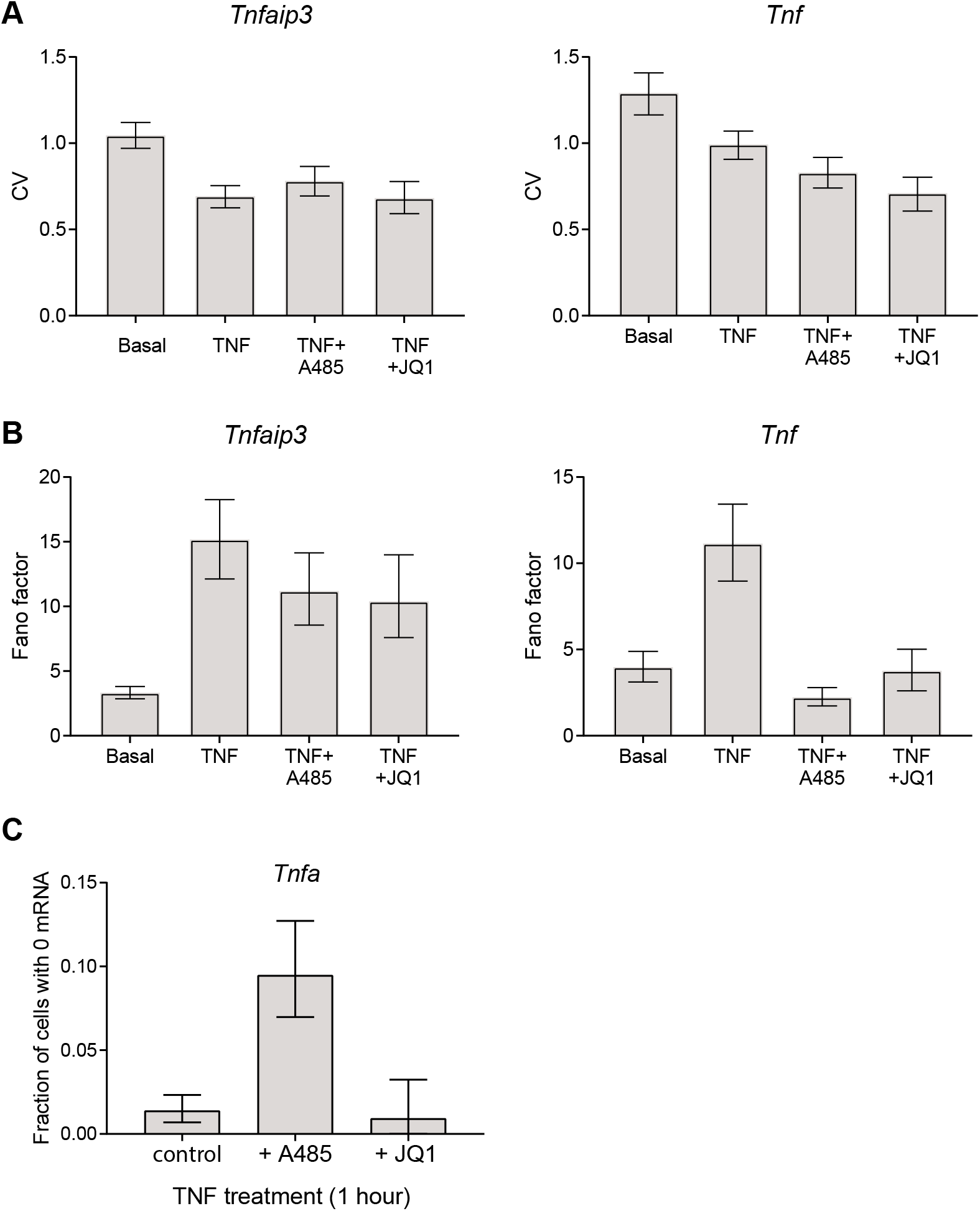
A-485 and JQ1 differentially affect noise and the fraction of non-responding cells in response to TNF treatment. **A-B** Bar graphs of coefficient of variation (CV) (B) and Fano factor (C) of TNF-stimulated smFISH distributions presented in Fig. 2B for the indicated genes at the indicated time points. Data are presented as mean and bootstrapped 95% CIs. **C** Fraction of cells with no Tnf transcripts after 1 hours of TNF stimulation in combination with the indicated inhibitors. Data are presented as mean +/- bootstrapped 95% CIs.

**Figure EV5 (related to Fig. 6).**
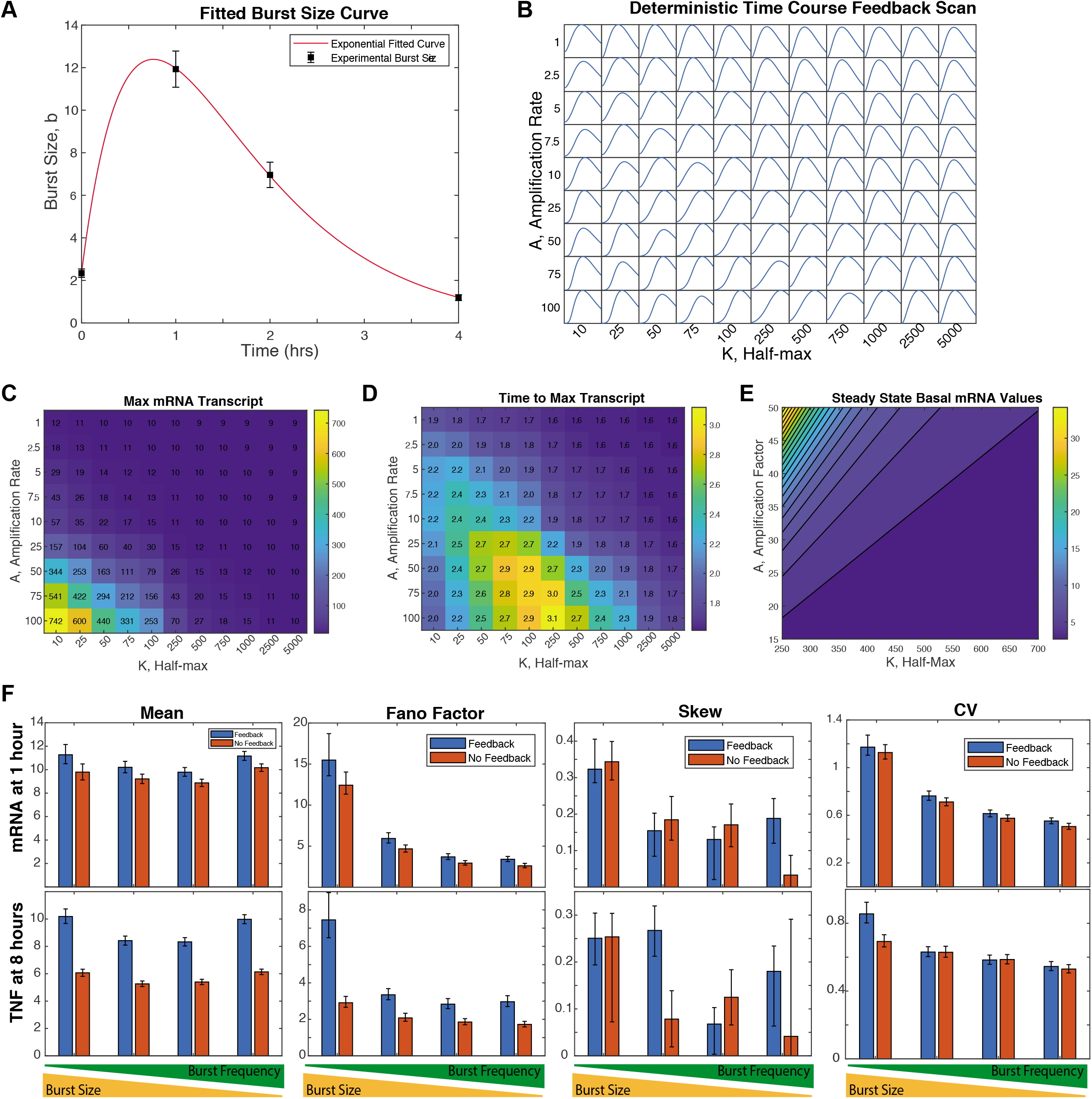
Parameter scans and fits for simulation of *Tnf* stochastic gene expression with positive feedback. **A** A time-dependent burst size function for Tnf mRNA production was fit to experimental data (Fig. 2). **B** Single-cell distributions of mRNA numbers are presented for varying values of the strength of TNF positive feedback (A) and the value of half-maximal strength (K). **C, D** Maximum mRNA transcript numbers (C) and time to reach maximum transcript number (D) for the parameter scan in (B). **E** Steady-state basal mRNA transcript levels for the parameter scan in (B). **F** Population mean, Fano factor, skew, and CV for mRNA (1 hour after TNF treatment) and protein (8 hours after TNF treatment) in simulated single cell populations for four different bursting parameter sets.

**Appendix Fig. 1 (related to Methods).**
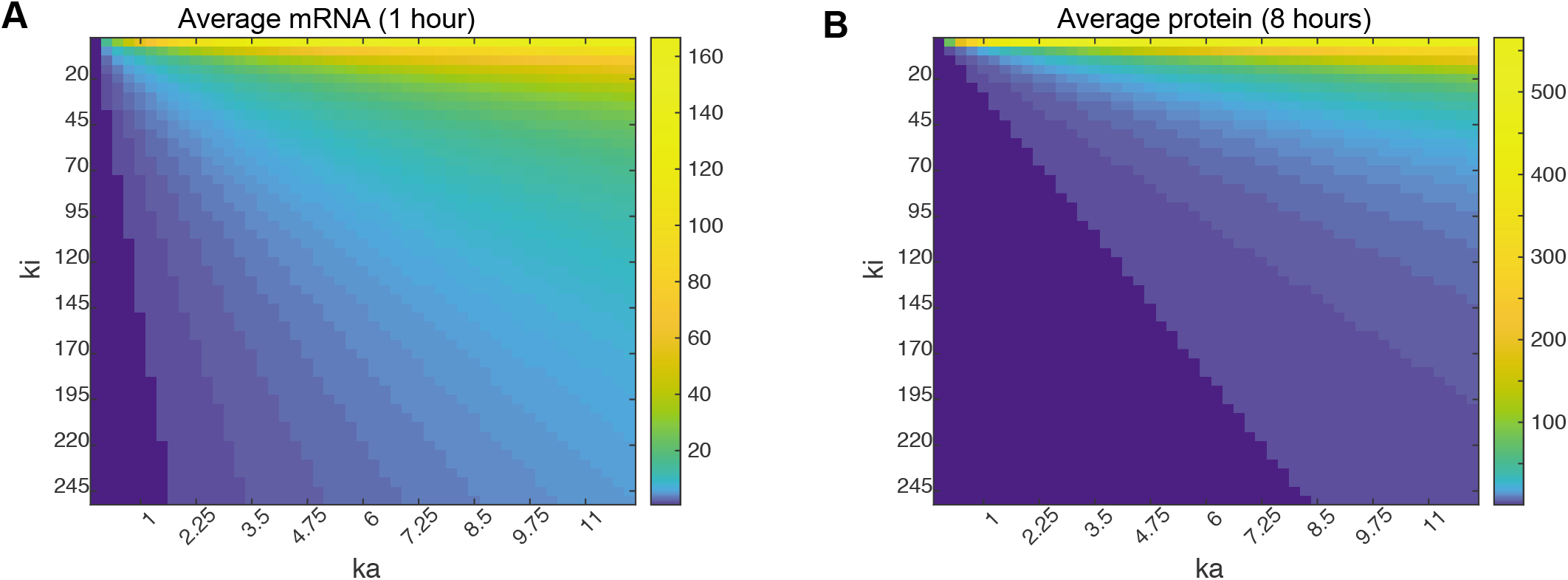
Parameter scans to find regions of mRNA expression. **A, B** Population averages from deterministic simulations of mRNA at 1 hour (A) and protein at 8 hours (B) after TNF treatment for the fitted kt value over a parameter scan of increasing values of ka and decreasing values of ki.

